# Antisense lncRNA transcription promotes A-to-I RNA editing via intermolecular dsRNA in breast cancer

**DOI:** 10.64898/2026.03.25.714158

**Authors:** Klaudia Samorowska, Elżbieta Wanowska, Michał Wojciech Szcześniak

## Abstract

A-to-I RNA editing, catalysed by ADAR enzymes, is the most prevalent post-transcriptional RNA modification in mammals, yet the regulatory inputs shaping cell-type-specific editomes remain incompletely understood. Here we characterise over 2.2 million unique A-to-I editing sites across MCF7 (ER+) and MDA-MB-231 (triple-negative) breast cancer cell lines and 117 patient tumours. MCF7 exhibited substantially more editing per sample, driven by higher ADAR1 expression and a shifted ADAR1/ADAR2 ratio that favoured broad intronic Alu editing in the luminal line versus site-selective synonymous coding editing in the aggressive line. Despite this divergence, approximately 2,500 sites were constitutively edited in both cell lines, defining a conserved core editome. We demonstrate that natural antisense lncRNA transcription constitutes an independent, additive pathway for editing through intermolecular dsRNA formation: editing density at sense-antisense overlaps reversed from depletion to enrichment as a function of balanced co-expression, antisense overlap increased editing probability without affecting density among edited genes, and a factorial analysis across all expressed genes established that inverted Alu pairs are the dominant editing substrate while antisense lncRNA transcription provides an independent contribution whose magnitude scales with overlap length and Alu content. Experimental validation at the *NDUFS1/NDUFS1-AS1* locus confirmed co-expression of sense and antisense transcripts, verified editing at computationally predicted positions by Sanger sequencing with genomic DNA controls, and demonstrated differential editing and expression between cell lines. Differentially edited genes included the oncogene *VOPP1* and fatty acid metabolism genes at antisense loci, linking epitranscriptomic regulation to the lipid metabolic phenotype of aggressive breast cancer. Our findings establish a two-tier model: a dominant ADAR1-driven programme targeting intramolecular Alu dsRNA, upon which an independent lncRNA antisense pathway is superimposed via intermolecular dsRNA, jointly producing subtype-specific editing landscapes that preserve a constitutive core but diverge in magnitude, functional distribution, and site selection.

## Introduction

Adenosine-to-Inosine (A-to-I) RNA editing is a widespread post-transcriptional modification in mammals catalysed by the adenosine deaminase acting on RNA (ADAR) family of enzymes. This process can influence multiple aspects of RNA metabolism, including stability (Vlachogiannis et al., 2021), splicing (Tang et al., 2020), translation efficiency and coding potential (Brachova et al., 2019), as well as modulate RNA structure and RNA–protein interactions (Delli Ponti et al., 2023). A-to-I editing occurs predominantly within non-coding regions of the transcriptome, particularly in repetitive Alu elements (Bazak et al., 2014a; Levanon et al., 2004). Although its biochemical basis is well understood, the sequence and structural determinants governing site-specific editing remain incompletely defined.

Dysregulation of A-to-I RNA editing has been increasingly linked to cancer. Altered ADAR expression and activity can reshape the transcriptome and promote tumorigenesis (Peng et al., 2018; Zhang et al., 2024). Elevated global editing levels, often driven by ADAR1 overexpression, have been associated with increased proliferation, immune evasion, and metastatic potential. At the molecular level, A-to-I editing results in deamination of adenosine to inosine which is interpreted as guanosine by the cellular machinery. Consequently, editing can recode codons, alter splice sites, and affect RNA structure as well as modulate interactions with proteins. Breast cancer (BC) is a heterogeneous disease comprising molecularly distinct subtypes with divergent clinical outcomes and regulatory programs. While RNA editing has been implicated in several cancers, its role in shaping differences between luminal and triple-negative breast cancer remains largely unexplored.

A-to-I editing is not limited to protein-coding transcripts but also extensively affects long non-coding RNAs (lncRNAs). Transcriptome-wide studies have revealed widespread editing within lncRNAs, particularly in double-stranded regions formed by repetitive elements such as Alu sequences (Silvestris et al., 2020). A well-characterised example is *NEAT1*, a nuclear-retained lncRNA enriched in paraspeckles, which contains multiple editing sites within Alu-derived dsRNA structures. ADAR1-mediated editing regulates *NEAT1* stability by modulating its interaction with RNA-binding proteins; inosine formation within AU-rich regions enhances binding of the stabilising factor AUF1, leading to increased *NEAT1* levels under inflammatory conditions. Consistently, ADAR1 overexpression elevates *NEAT1* abundance, whereas its depletion reduces *NEAT1* expression and impairs its induction. These observations highlight a direct link between RNA editing and lncRNA stability (Vlachogiannis et al., 2021).

Beyond serving as substrates, lncRNAs can actively shape RNA editing landscapes. For instance, the snoRNA-derived lncRNA *LNC-SNO49AB* promotes ADAR1 homodimerisation, thereby enhancing global editing activity and influencing downstream gene expression (Huang et al., 2022). Additionally, natural antisense lncRNAs can regulate editing at specific loci through RNA-RNA interactions. The prostate cancer-associated lncRNA *PCA3* forms intermolecular duplexes with its sense counterpart *PRUNE2*, promoting A-to-I editing and consequent downregulation of *PRUNE2* (Salameh et al., 2015). Another example is the antisense lncRNA *UGGT1-AS1*, which regulates expression of its cognate sense gene *UGGT1* in breast cancer cells. Building on this observation as well as high expected frequencies of such events in the scale of full transcriptomes (Szcześniak and Makałowska, 2016), we previously proposed that *UGGT1*-*AS1* may act as a triggering factor for A-to-I RNA editing in the sense transcript, suggesting a role for antisense transcription in generating editing-competent dsRNA structures (Samorowska et al., 2025). Together, these findings indicate that lncRNAs influence RNA editing both by modulating ADAR activity and by shaping the availability of double-stranded RNA substrates. A particularly relevant class are natural antisense transcripts (NATs), which overlap protein-coding genes and can form intermolecular dsRNA duplexes with their sense counterparts. Because dsRNA constitutes the primary substrate for ADAR enzymes, antisense transcription represents a plausible yet underexplored mechanism for directing RNA editing. We therefore hypothesise that sense–antisense pairing promotes A-to-I editing through the formation of intermolecular dsRNA structures.

Here, we investigated the determinants of RNA editing in breast cancer by analysing transcriptome-wide A-to-I editing landscapes across luminal (MCF7) and triple-negative (MDA-MB-231) breast cancer cell lines, as well as primary tumour samples using publicly available data. We identify distinct editing programs associated with molecular subtype, driven in part by differences in ADAR expression and editing substrate availability. Furthermore, we demonstrate that antisense lncRNA transcription constitutes an independent and additive mechanism contributing to RNA editing through the formation of intermolecular dsRNA. Using integrative computational analyses and experimental validation, we show that while intramolecular Alu-derived dsRNA remains the dominant substrate for ADAR activity, antisense transcription provides an additional regulatory layer that shapes transcript-specific editing patterns. Together, our findings support a two-tier model of RNA editing regulation, in which a core ADAR1-driven program is complemented by lncRNA-mediated mechanisms, contributing to subtype-specific epitranscriptomic landscapes in breast cancer.

## Results

### Global A-to-I editing landscape and Alu validation

To investigate A-to-I RNA editing across breast cancer subtypes, we identified editing sites using the SPRINT toolkit in 45 MCF7 (ER+) and 79 MDA-MB-231 (TNBC) RNA-seq samples, followed by annotation with Ensembl Variant Effect Predictor (VEP, GRCh38). After removing known genomic variants (ClinVar-annotated, gnomAD ≥0.1% allele frequency, and dbSNP entries) and retaining only A→I-consistent changes (A>G and T>C), we identified 2,232,436 unique editing sites: 1,387,735 variants in MCF7 and 1,217,066 in MDA-MB-231. The two cell lines shared 372,365 editing sites, with 1,015,370 sites unique to MCF7 and 844,701 unique to MDA-MB-231 (**Figure 1A**). Consistent with ADAR-mediated editing, 82.2% of all sites localised within Alu repetitive elements, which form intramolecular dsRNA structures preferentially targeted by ADAR enzymes. The Alu overlap fraction was slightly but significantly higher in MCF7 than in MDA-MB-231 (84.8% vs 82.0% of unique sites, Fisher’s exact p < 2.2 × 10⁻¹⁶; per-sample median 88.1% vs 86.8%, Mann-Whitney p = 0.047; **Figure 1B,C**), foreshadowing the ADAR expression differences described below.

**Figure 1.**
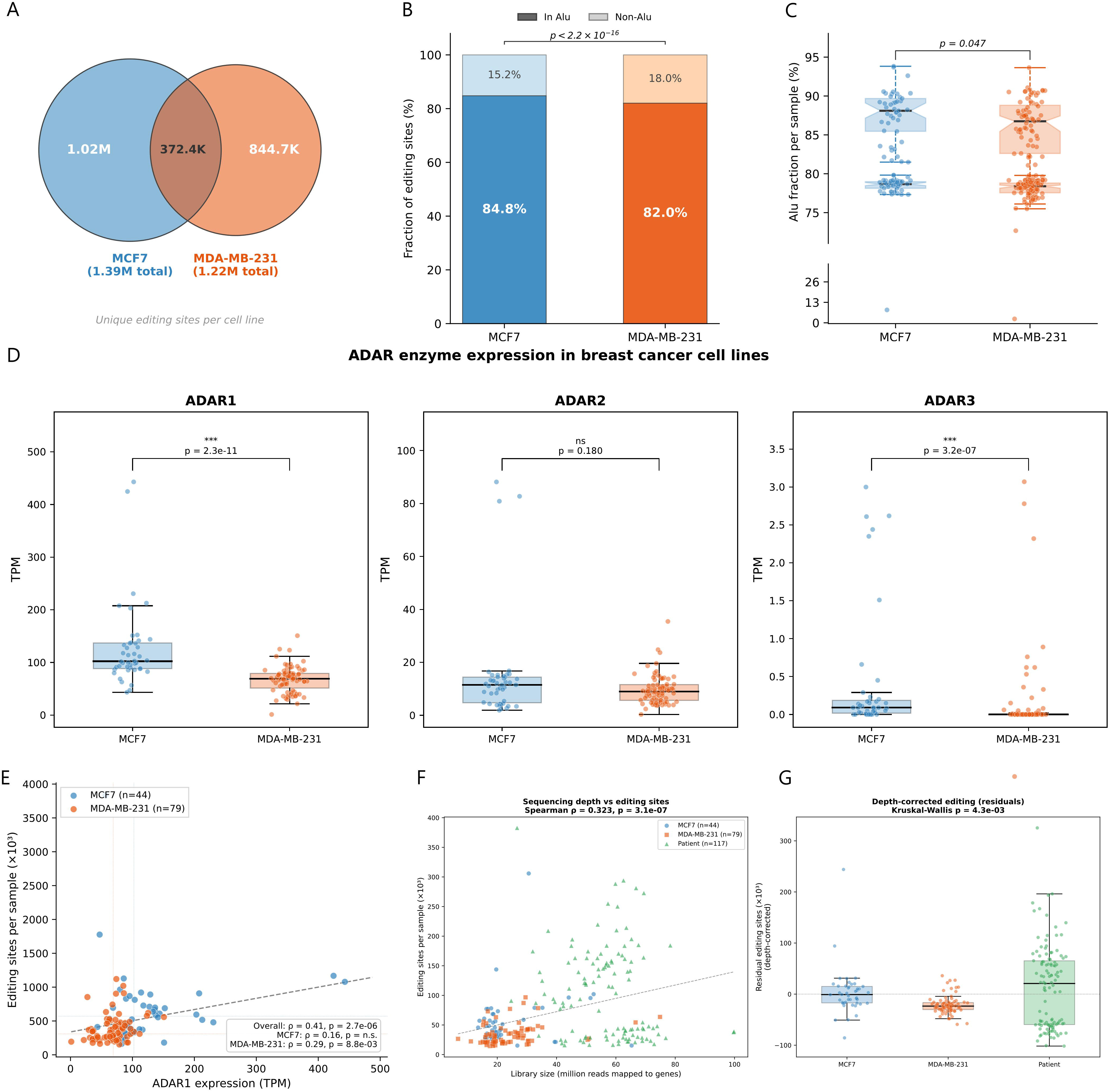
**(A)** Proportional Venn diagram showing the overlap of unique editing sites between MCF7 (ER+, blue) and MDA-MB-231 (TNBC, orange). MCF7 harbored 1,387,735 unique sites and MDA-MB-231 harbored 1,217,066, with 372,365 sites shared between cell lines. Circle areas are proportional to set sizes. **(B)** Fraction of unique editing sites overlapping Alu repetitive elements per cell line. Dark bars indicate sites within Alu elements; light bars indicate non-Alu sites. MCF7 showed a significantly higher Alu overlap fraction than MDA-MB-231 (84.8% vs 82.0%; Fisher’s exact test p < 2.2 × 10⁻¹⁶). The high overall Alu fraction (82.2% across both lines) confirms ADAR-mediated editing as the dominant mechanism. **(C)** Per-sample Alu fraction distributions. Each point represents one RNA-seq sample. Boxplots show median (black line), interquartile range (shaded box), and whiskers extending to 1.5× IQR. MCF7 samples showed a modestly but significantly higher per-sample Alu fraction than MDA-MB-231 (median 88.1% vs 86.8%; Mann-Whitney U test p = 0.047). The broken y-axis accommodates outlier samples with anomalously low Alu fractions. **(D)** Expression of ADAR family enzymes (ADAR1, ADAR2, ADAR3) in MCF7 (blue, n=45) and MDA-MB-231 (orange, n=79) samples, quantified as transcripts per million (TPM) from RSEM. **(E)** Relationship between ADAR1 expression and per-sample editing site count. Each point represents one RNA-seq sample (44 MCF7, 79 MDA-MB-231). The dashed line indicates the overall linear trend. Dotted crosshairs mark the median. **(F)** Scatter plot of library size (million reads mapped to genes) versus unique editing sites per sample for MCF7 (blue circles, n=44), MDA-MB-231 (orange squares, n=79), and patient tumors (green triangles, n=117). The dashed line indicates the overall linear trend (Spearman ρ = 0.32, p = 3.1 × 10⁻⁷). Library sizes were comparable between cell lines (median 19.5 × 10⁶ vs 21.8 × 10⁶; Mann-Whitney p = 0.28), whereas patient samples were sequenced considerably more deeply (median 58.5 × 10⁶; p = 5.7 × 10⁻¹⁹ vs cell lines). Within MCF7, editing was uncorrelated with depth (ρ = 0.05, p = 0.77); MDA-MB-231 showed a modest positive association (ρ = 0.36, p = 0.001); patients showed no depth–editing correlation (ρ = −0.13, p = 0.16). **(G)** Residual editing site counts after linear regression on library size across all 240 samples (R² = 0.115), shown as boxplots with individual sample points. After depth correction, cohort differences persisted (Kruskal-Wallis p = 4.3 × 10⁻³): MCF7 showed a mean residual of +3,278 sites, MDA-MB-231 showed −21,069, and patients showed +12,993 above the depth-predicted baseline. The horizontal dotted line indicates zero residual.

### Differential ADAR enzyme expression underlies subtype-specific editing

To investigate the molecular basis of the editing differences between cell lines, we compared expression of ADAR family enzymes from the same RNA-seq datasets (RSEM-derived TPM). ADAR1 (ADAR), the principal editor of Alu-containing dsRNA, was significantly overexpressed in MCF7 relative to MDA-MB-231 (median 102.3 vs 69.0 TPM; Mann-Whitney p = 2.3 × 10⁻¹¹; Cohen’s d = 1.04; **Figure 1D**), consistent with the 1.6-fold higher per-sample editing site count in MCF7 (median 46,990 vs 28,671). In contrast, ADAR2 (ADARB1), which preferentially targets specific coding sites, showed comparable expression (median 11.5 vs 9.0 TPM; p = 0.18), while the catalytically inactive ADAR3 (ADARB2) was minimally expressed in both lines (median < 0.1 TPM). The ADAR1-to-ADAR2 expression ratio was significantly shifted toward ADAR1 in MCF7 (median 9.4 vs 7.4; Mann-Whitney p = 2.0 × 10⁻³), suggesting a cell-line-specific enzyme stoichiometry that favors broad Alu-targeted editing in the luminal line. Additionally, ADAR1 expression correlated positively with the per sample editing site count across the combined cohort (Spearman ρ = 0.41, p = 2.7 × 10⁻⁶; **Figure 1E**), with the association driven primarily by between-group differences: MCF7 samples clustered at higher ADAR1 levels (median 102.3 TPM) and higher editing site counts (median 46,990), while MDA-MB-231 samples clustered at lower values (69.0 TPM, 28,671 sites).

### Editing site counts are independent of sequencing depth

To rule out technical confounders, we assessed the relationship between sequencing depth and editing site detection across all three cohorts. Library sizes were comparable between cell lines (median 19.5 × 10⁶ vs 21.8 × 10⁶ mapped reads for MCF7 and MDA-MB-231; Mann-Whitney p = 0.28), whereas patient tumour samples were sequenced considerably more deeply (median 58.5 × 10⁶; Mann-Whitney p = 5.7 × 10⁻¹⁹ vs cell lines). Across all 240 samples, sequencing depth showed a modest overall correlation with editing site count (Spearman ρ = 0.32, p = 3.1 × 10⁻⁷; R² = 0.115; **Figure 1F**), driven largely by the between-cohort difference in library size. Within individual cohorts, the association was absent in MCF7 (ρ = 0.05, p = 0.77) and in patients (ρ = −0.13, p = 0.16), whereas MDA-MB-231 showed a modest positive association (ρ = 0.36, p = 0.001), indicating that some editing heterogeneity in this aggressive line is amplified at greater depth. After regressing out the depth effect across all 240 samples, cohort differences persisted (Kruskal-Wallis p = 4.3 × 10⁻³): MCF7 showed a mean residual of +3,278 sites, MDA-MB-231 showed −21,069, and patients showed +12,993 above the depth-predicted baseline (**Figure 1G**), confirming that the editing differences between cohorts reflect genuine biological variation rather than sequencing artefacts.

### Subtype-specific functional distribution of editing sites

The overall functional distribution of editing sites differed markedly between cell lines (χ² = 5,345, df = 9, p < 2.2 × 10⁻¹⁶; **Figure 2A**). Intronic editing dominated both lines but was proportionally enriched in MCF7 (72.6% vs 69.9%; OR = 1.14, Fisher’s exact p < 2.2 × 10⁻¹⁶), consistent with its higher ADAR1/ADAR2 ratio and Alu fraction. Conversely, MDA-MB-231 showed proportional enrichment in UTR (9.4% vs 8.5%; OR = 0.90, p = 4.1 × 10⁻¹⁴²), intergenic (8.9% vs 8.2%), and upstream/downstream regions, suggesting a redistribution of editing activity toward functional non-coding targets in the aggressive line.

**Figure 2.**
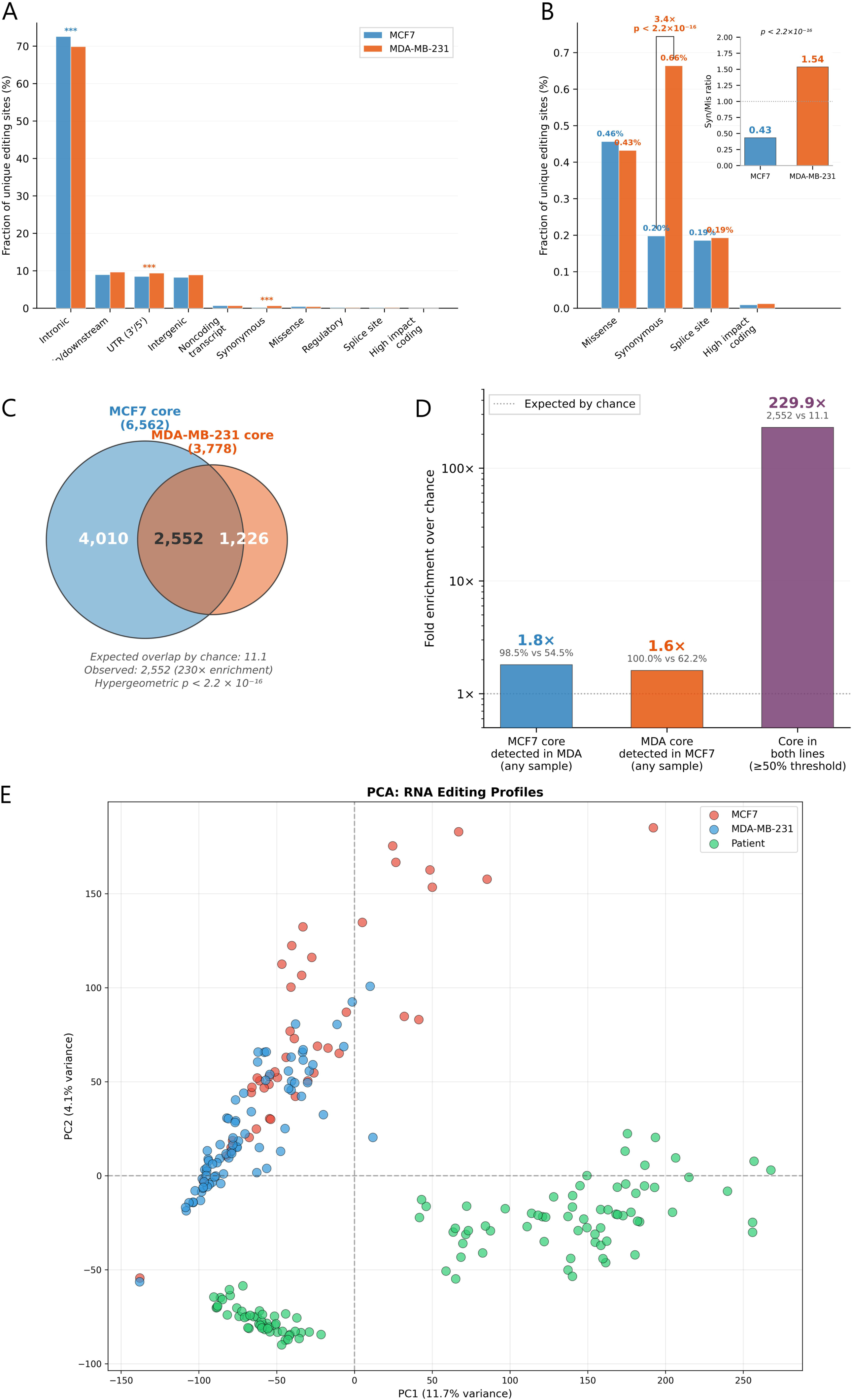
Subtype-specific functional redistribution of editing sites. **(A)** Grouped bar chart showing the percentage of unique editing sites in each functional category for MCF7 (blue, 1,387,735 sites) and MDA-MB-231 (orange, 1,217,066 sites). Categories are ordered by abundance. Intronic editing dominated both lines but was proportionally enriched in MCF7 (72.6% vs 69.9%; OR = 1.14, p < 2.2 × 10⁻¹⁶), while UTR editing was enriched in MDA-MB-231 (9.4% vs 8.5%; OR = 0.90, p = 4.1 × 10⁻¹⁴²). The overall functional distribution differed significantly between cell lines (χ² = 5,345, df = 9, p < 2.2 × 10⁻¹⁶). Asterisks indicate significance after Bonferroni correction (*** p < 4.55 × 10⁻³). **(B)** A conserved core editome persists across divergent editing landscapes. Zoomed view of coding and regulatory categories. The most striking difference was in synonymous editing: MDA-MB-231 exhibited 3.4-fold higher proportional synonymous editing than MCF7 (0.66% vs 0.20%; OR = 0.30, p < 2.2 × 10⁻¹⁶), while missense (0.46% vs 0.43%) and splice site editing (0.19% both) were comparable. Inset: synonymous-to-missense ratio was inverted between cell lines (MCF7 = 0.43, MDA-MB-231 = 1.54; Fisher’s exact p < 2.2 × 10⁻¹⁶). **(C)** Proportional Venn diagram of core editome sites (present in ≥50% of samples within each cell line). MCF7 defined 6,562 core sites (≥23/45 samples) and MDA-MB-231 defined 3,778 core sites (≥40/79 samples), with 2,552 sites independently classified as core in both lines. Circle areas are proportional to set sizes. **(D)** Fold enrichment of observed over expected overlap for three progressively stringent tests of core editome sharing. Left: 98.5% of MCF7 core sites were detected in at least one MDA-MB-231 sample, compared to 54.5% expected by chance (1.8-fold). Center: 100.0% of MDA-MB-231 core sites were detected in MCF7, versus 62.2% expected (1.6-fold). Right: 2,552 sites were core in both cell lines, versus 11.1 expected (230-fold). The dotted line indicates the chance baseline (1×). **(E)** Patient tumor editomes cluster distinctly from cell lines but preserve the functional sharing hierarchy. Principal component analysis of per-sample editing site profiles across MCF7 (red, n=45), MDA-MB-231 (blue, n=79), and patient tumors (green, n=117). Patient samples separate into two subclusters (circles vs squares) that correlate with sequencing batch dates (62% of subcluster 0 sequenced Feb 22, 2016; 89.6% of subcluster 1 sequenced Jan 27, 2016), indicating a technical batch effect in the clinical cohort. Despite this confound, patients cluster entirely separately from both cell lines, demonstrating that in vitro culture does not fully recapitulate tumor editing complexity.

The most striking difference was in synonymous coding sites: MDA-MB-231 exhibited 0.66% synonymous editing sites versus 0.20% in MCF7, a 3.4-fold proportional enrichment (OR = 0.30, Fisher’s exact p < 2.2 × 10⁻¹⁶; **Figure 2B**). This was not a general coding-region effect: missense variants showed a slight MCF7 enrichment (0.46% vs 0.43%; OR = 1.06, p = 3.3 × 10⁻³), and splice site editing was equivalent between cell lines (0.19% in both; p = 0.20). Within coding regions, the synonymous-to-missense ratio crossed unity in opposite directions: 0.43 in MCF7 (missense-dominated) versus 1.54 in MDA-MB-231 (synonymous-dominated; Fisher’s exact p < 2.2 × 10⁻¹⁶; **Figure 2B inset**), indicating that MCF7 editing preferentially recodes amino acids while MDA-MB-231 editing targets synonymous positions, potentially influencing translation kinetics or RNA secondary structure without disrupting protein sequence.

### A conserved core editome persists across divergent editing landscapes

Despite fundamental differences in editing magnitude and functional distribution, MCF7 and MDA-MB-231 maintained a shared core of highly reproducible editing sites. The majority of editing sites showed low reproducibility: 75.0% of MCF7 variants and 73.4% of MDA-MB-231 variants were singletons detected in only one sample, with mean intra-cell-line sharing rates of 4.7% (MCF7) and 3.2% (MDA-MB-231). However, defining common variants as those present in ≥50% of samples revealed a substantial core editome: 6,562 sites in MCF7 (≥23 of 45 samples) and 3,778 sites in MDA-MB-231 (≥40 of 79 samples). Nearly all core editome sites were detected in the other cell line: 98.5% of MCF7’s common variants (6,462/6,562) and 100.0% of MDA-MB-231’s common variants (3,777/3,778), significantly exceeding the 54.5% and 62.2% expected by chance (hypergeometric p < 2.2 × 10⁻¹⁶ for both; **Figure 2C**). More stringently, 2,552 sites were independently classified as core in both cell lines - a 230-fold enrichment over the 11.1 expected if core status were randomly distributed (hypergeometric p < 2.2 × 10⁻¹⁶; **Figure 2D**). This shared constitutive editome suggests a deterministic ADAR-dependent program that maintains editing at several thousand key sites, upon which subtype-specific variation is layered.

### Patient tumor editomes confirm cell-line patterns but reveal depth-dependent inflation

Analysis of 117 dbGaP patient tumour samples revealed 4,672,341 unique RNA editing sites with a median of 93,750 sites per patient, substantially exceeding cell-line counts - consistent with both the deeper sequencing and the greater biological heterogeneity of primary tumours. Principal component analysis of per-sample editing profiles revealed that patients clustered distinctly from both cell lines (**Figure 2E**), consistent with tumor cellular heterogeneity not recapitulated in culture. Notably, patient samples separated into two subclusters that correlated with sequencing dates (62% of subcluster 0 from Feb 22, 2016; 89.6% of subcluster 1 from Jan 27, 2016), highlighting technical batch effects as a confound in clinical editome analyses. Despite this variation, the patient core editome comprised 5,177 sites, intermediate between MCF7 (6,562) and MDA-MB-231 (3,778), with 55.5% localizing to UTRs.

Category-specific mean sharing rates preserved the functional hierarchy observed in cell lines across all three cohorts - UTR (9.4%, 6.6%, 6.1% for MCF7, MDA-MB-231, and patients, respectively) > splice site (6.1%, 4.0%, 4.1%) > noncoding transcript (6.0%, 3.9%, 3.8%) > regulatory > intronic > intergenic (**Supplementary Figure 1**) - providing evidence for conserved, functionally driven RNA editing programs in breast cancer that persist from cultured cells to primary tumors despite batch-related noise.

### Antisense lncRNA transcription promotes A-to-I editing through dsRNA formation

#### Editing density is enriched in expressed sense–antisense overlapping regions

To investigate whether lncRNA:sense transcript interactions could trigger A-to-I editing through ADAR, we mapped 2,232,436 unique editing sites onto the human genome annotation. The total non-redundant length of annotated genes was 2,337,576,590 nucleotides, of which 272,317,835 nucleotides (11.6%) represented sense–antisense overlapping regions on a single genomic strand. At the genome-wide level, editing density was significantly lower in overlapping than in non-overlapping regions (0.056% vs 0.085%; 0.66-fold; χ² = 44,874, p < 2.2 × 10⁻¹⁶; **Figure 3A**), indicating that genomic overlap per se does not promote editing when transcriptional activity is not considered. This pattern reversed when we restricted the analysis to actively transcribed genes (≥5 TPM), under which overlapping regions showed significantly higher editing density (0.153% vs 0.119%; 1.28-fold; χ² = 7,568, p < 2.2 × 10⁻¹⁶).

**Figure 3.**
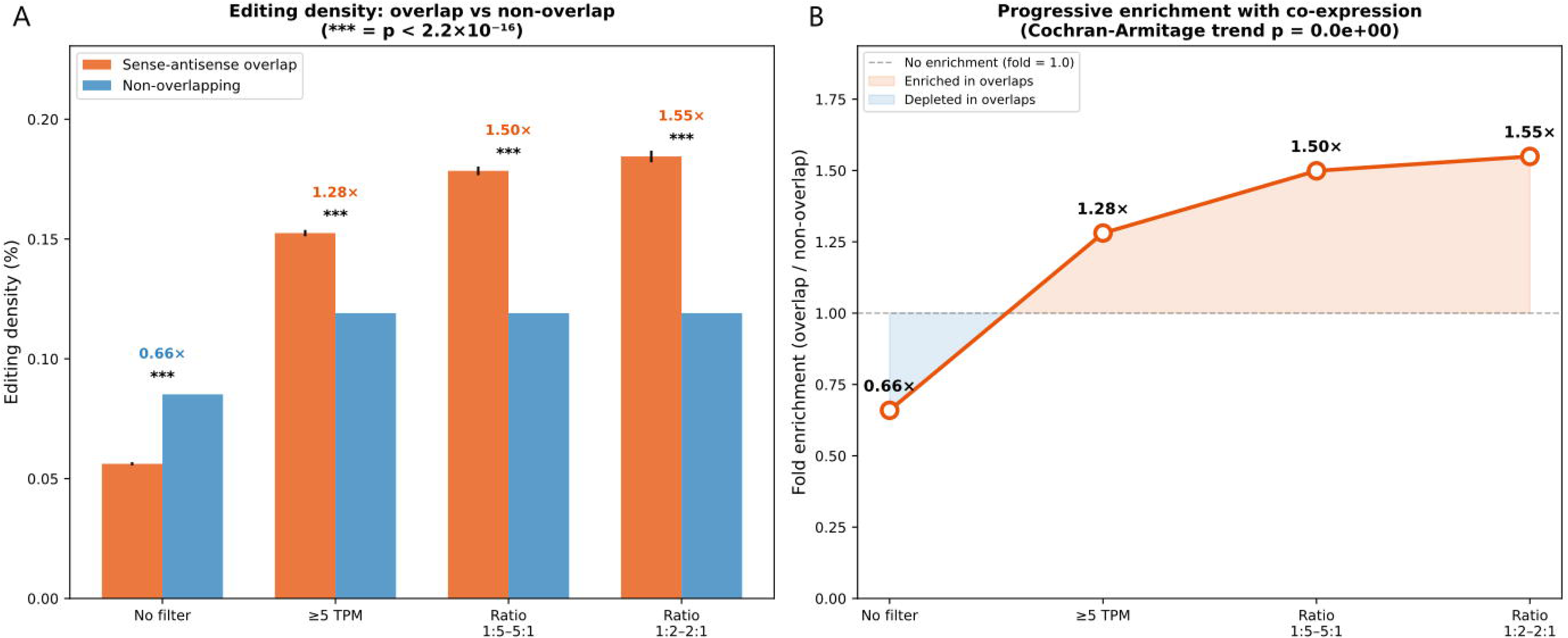
Antisense lncRNA transcription promotes A-to-I editing through dsRNA formation. Editing density is enriched in expressed sense–antisense overlapping regions. **(A)** Editing density (% of nucleotides with editing sites) in sense–antisense overlapping regions (orange) versus non-overlapping regions (blue) across four progressively stringent expression filters. Without expression filtering, overlapping regions were significantly depleted for editing sites relative to non-overlapping regions (0.056% vs 0.085%; 0.66-fold; χ² = 44,874, p < 2.2 × 10⁻¹⁶). This pattern reversed upon filtering for actively transcribed genes (≥5 TPM: 1.28-fold; expression ratio 1:5–5:1: 1.50-fold; ratio 1:2–2:1: 1.55-fold). All comparisons p < 2.2 × 10⁻¹⁶ (***). Error bars indicate 95% Wilson confidence intervals. **(B)** Fold enrichment of editing density in overlapping versus non-overlapping regions as a function of expression filter stringency. The Cochran–Armitage trend test confirmed a monotonically significant increase in editing density with increasingly balanced co-expression (Z = 39.7, p < 2.2 × 10⁻¹⁶). The dashed line indicates no enrichment (fold = 1.0); shading indicates enrichment (orange, above) or depletion (blue, below) relative to non-overlapping regions.

We further refined this analysis by imposing balanced expression requirements. Restricting to overlaps with expression ratios between 1:5 and 5:1 yielded 0.179% density (1.50-fold; χ² = 11,223, p < 2.2 × 10⁻¹⁶). Tightening to 1:2–2:1 yielded 0.185% density (1.55-fold; χ² = 6,767, p < 2.2 × 10⁻¹⁶). A Cochran–Armitage trend test confirmed the monotonic increase (Z = 39.7, p < 2.2 × 10⁻¹⁶; **Figure 3B**). The reversal from depletion (0.66-fold) to progressive enrichment (1.28 → 1.50 → 1.55-fold) with increasingly balanced co-expression demonstrates that the editing enrichment at sense-antisense overlaps is specifically dependent on active bidirectional transcription.

#### Antisense expression correlates positively with editing efficiency

We next tested whether antisense transcript abundance quantitatively correlates with editing efficiency. We calculated Spearman correlations between the antisense:sense expression ratio and the editing ratio across all expressed gene pairs (≥10 samples, both partners ≥10 TPM), yielding 332 testable lncRNA antisense sites and 1,947 protein-coding antisense sites.

Across all tested sites, correlation coefficients were significantly skewed toward positive values for both biotypes (**Supplementary Figure 2,** left panels). Among 332 lncRNA sites, 182 (54.8%) exhibited positive correlations (binomial p = 0.019; mean ρ = 0.098, 95% CI [0.054, 0.141]; t-test p = 1.4 × 10⁻⁵). Among 1,947 protein-coding sites, 1,053 (54.1%) were positive (binomial p = 2.2 × 10⁻⁵; mean ρ = 0.059, 95% CI [0.042, 0.077]; t-test p = 2.0 × 10⁻¹¹). The proportion did not differ between biotypes (Fisher’s exact p = 0.59), confirming a general architectural feature independent of transcript type.

After Benjamini–Hochberg correction, 54 lncRNA sites (16.3%) and 222 protein-coding sites (11.4%) remained significant at FDR < 0.05 - substantially exceeding the 16.6 and 97.4 false positives expected by chance (**Supplementary Figure 2**, centre and right panels). The positive directional bias strengthened in the FDR-corrected subsets: 70.4% of significant lncRNA correlations (38/54; binomial p = 0.004) and 65.3% of protein-coding correlations (145/222; binomial p = 5.9 × 10⁻⁶) were positive.

### Gene-level evidence for antisense-dependent editing

#### Antisense lncRNA transcription increases the probability but not the density of gene-level editing

Among 12,653 expressed protein-coding genes (≥1 TPM, ≥1 kb), those overlapping an antisense lncRNA were significantly more likely to harbor editing events: 25.8% versus 17.3% in MCF7 (OR = 1.66, Fisher’s exact p = 5.2 × 10⁻¹⁷), 15.6% versus 11.6% in MDA-MB-231 (OR = 1.41, p = 3.6 × 10⁻⁶), and 27.7% versus 19.2% in the combined editome (OR = 1.61, p = 7.5 × 10⁻¹⁶; **Figure 4A**). This effect persisted within gene-length quartiles and expression bins. However, among edited genes, per-sample editing density was comparable regardless of antisense lncRNA status (MCF7: median 0.160 vs 0.140 sites/kb, p = 0.60; MDA-MB-231: 0.153 vs 0.164, p = 0.92; **Figure 4B**), indicating that the presence of an annotated antisense lncRNA governs the probability of editing rather than its density. This binary classification does not capture quantitative variation in antisense expression levels; the positive correlation between antisense:sense expression ratio and editing efficiency described above indicates that among actively co-expressed pairs, antisense transcript abundance does modulate editing output in a dose-dependent manner. The two observations are reconciled by the modest effect size of the quantitative association (mean ρ = 0.098) and the dominance of intramolecular Alu-pair substrates in determining overall editing density, such that the antisense contribution is detectable as a continuous within-gene signal but not as a shift in median density across the binary gene grouping.

**Figure 4.**
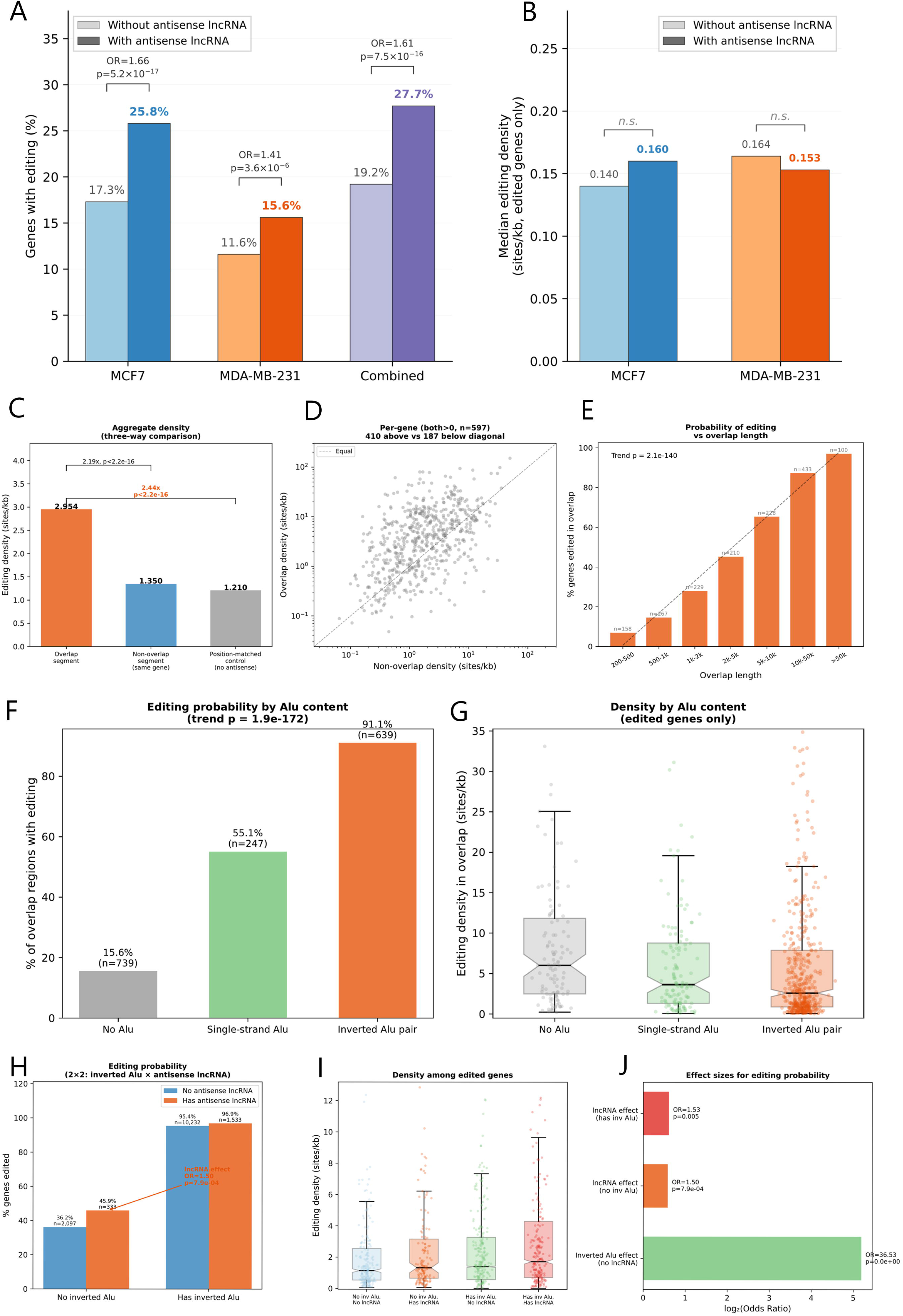
Gene-level evidence for antisense-dependent editing. Antisense lncRNA transcription increases the probability but not the density of gene-level editing**. (A)** Percentage of expressed protein-coding genes (n = 12,653) harbouring editing events, stratified by antisense lncRNA status. Brackets show odds ratios and Fisher’s exact p-values. **(B)** Median per-sample editing density among edited genes only, showing no significant difference by antisense lncRNA status in either cell line. **(C)** Editing localises to the region of sense–antisense interaction. Aggregate editing density in the overlap segment, the non-overlap segment of the same gene, and position-matched control segments from genes lacking antisense lncRNA. Brackets show fold differences and p-values. **(D)** Per-gene scatter comparing overlap versus non-overlap density (log-scaled; dashed line = equal density). **(E)** Editing probability scales with antisense overlap length. Percentage of genes edited in each overlap length bin (n shown above bars; Cochran–Armitage trend line dashed). **(F)** Inverted Alu elements mediate editing at sense–antisense overlaps. Editing probability by Alu configuration within 1,625 overlap regions. **(G)** Editing density among edited overlaps stratified by Alu configuration; notched boxes show 95% CI for the median. **(H)** Intramolecular and intermolecular dsRNA operate as complementary editing pathways. Editing probability in a 2×2 factorial design crossing inverted Alu status with antisense lncRNA status across 14,195 genes. **(I)** Editing density among edited genes in each category; notched boxes show 95% CI for the median. **(J)** Effect sizes (log₂ odds ratio) for the three independent comparisons.

#### Editing localises to the region of sense-antisense interaction

For 1,411 protein-coding genes with partial antisense lncRNA overlap, we compared editing density in the overlap segment versus position-matched control segments from 11,297 genes lacking antisense lncRNA (median overlap position: 10–55% of gene body from the 5′ end). Overlap segments exhibited 2.44-fold higher editing density than position-matched controls (2.95 vs 1.21 sites/kb; χ² = 36,935, p < 2.2 × 10⁻¹⁶) and 2.19-fold higher than non-overlap segments of the same genes (1.35 sites/kb; χ² = 23,726, p < 2.2 × 10⁻¹⁶; **Figure 4C**). Since this comparison controls for positional context within the gene body, the enrichment is attributable to the antisense interaction itself rather than to 5′/3′ compositional biases. Per-gene paired comparisons confirmed that overlap density exceeded non-overlap density in the majority of individual genes (410 above vs 187 below the diagonal among 597 genes with editing in both compartments; **Figure 4D**). Antisense overlap start positions were concentrated near the 3′ end of the gene body (**Supplementary Figure 3A**), consistent with typical antisense lncRNA architecture.

#### Editing probability scales with overlap length

Among 1,625 protein-coding genes with antisense lncRNA overlaps, editing probability increased monotonically with overlap length: from 7.0% for 200-500 bp overlaps to 97.0% for overlaps exceeding 50 kb (Cochran-Armitage Z = 25.2, p = 2.1 × 10⁻¹⁴⁰; **Figure 4E**). After controlling for gene length, overlap length remained strongly associated with editing probability (partial Spearman ρ = 0.67, p = 1.7 × 10⁻¹⁹⁸) but not with editing density (ρ = −0.006, p = 0.83; **Supplementary Figure 3B**). The overlap fraction relative to gene length also predicted editing (Spearman ρ = 0.17, p = 8.8 × 10⁻¹¹; **Supplementary Figure 3C**), with genes having 40–95% overlap reaching 90.9% editing probability compared to 21.2% for those with less than 5%.

#### Inverted Alu elements mediate editing at sense–antisense overlaps

Within overlap regions, editing probability followed a steep gradient with Alu element content: 15.6% of Alu-free overlaps were edited, compared to 55.1% with Alu on one genomic strand and 91.1% with Alu on both strands (Cochran–Armitage Z = 28.0, p = 1.9 × 10⁻¹⁷²; inverted vs no Alu: OR = 55.4, p = 2.8 × 10⁻¹⁹⁴; **Figure 4F**). The number of Alu elements correlated strongly with editing density across all 1,625 overlap regions (Spearman ρ = 0.61, p = 1.1 × 10⁻¹⁶³; ρ = 0.27, p = 3.1 × 10⁻¹⁶ among 833 edited genes only; **Figure 4G, Supplementary Figure 3D**), and 89.0% of editing sites in Alu-containing overlaps localised within the Alu elements themselves (per-gene median 96.7%). Since both sense and antisense transcripts traverse the overlap region, even a single Alu insertion provides complementary sequence for intermolecular dsRNA formation between the two transcripts.

#### Intramolecular and intermolecular dsRNA operate as complementary editing pathways

To dissect the relative contributions of intramolecular dsRNA (inverted Alu pairs within a single transcript) and intermolecular dsRNA (formed between sense and antisense transcripts), we classified all 14,195 protein-coding genes by two independent properties. The majority (83%) harbored their own inverted Alu pairs, which was the dominant determinant of editing: 95.4% of such genes were edited compared to 36.2% without (OR = 36.5, p < 2.2 × 10⁻¹⁶; **Figure 4H**). Antisense lncRNA transcription provided a significant additional contribution: among genes lacking inverted Alu pairs, those with an antisense lncRNA were 1.27-fold more likely to be edited (45.9% vs 36.2%; OR = 1.50, p = 7.9 × 10⁻⁴). The lncRNA effect was virtually identical among genes with inverted Alu (OR = 1.53, p = 5.0 × 10⁻³), with no interaction between the two mechanisms (p = 0.93; **Figure 4I**). These findings establish a hierarchical model: inverted Alu elements constitute the primary editing substrate (OR = 36.5), while antisense lncRNA transcription provides an independent, additive pathway (OR ≈ 1.5) via intermolecular dsRNA (**Figure 4J)**, with the magnitude of this effect scaling with overlap length and Alu content.

### Subtype-specific gene editing programs and pathway enrichment

#### Gene-level editing density reveals subtype-specific programs

To identify genes with subtype-specific editing patterns, we computed per-sample editing density (sites/kb) for each gene in each cell line and compared the distributions using Mann-Whitney tests with Benjamini–Hochberg correction. Among 9,538 genes tested (edited in ≥5 samples of both cell lines), 931 showed significantly different editing density (FDR < 0.05), with a strong asymmetry: 373 genes were MCF7-enriched (≥2-fold higher median density) versus only 38 MDA-enriched (**Figure 5A**), consistent with MCF7’s higher ADAR1 expression driving broader editing. The most-edited genes in both cell lines were predominantly lncRNAs (all top 17 in MCF7, top 7 in MDA), reaching densities of 20–56 sites/kb - consistent with lncRNAs forming extensive dsRNA structures targeted by ADAR. MCF7-enriched genes were over-represented for lncRNAs (27.9% vs 17.0% background), while MDA-enriched genes were predominantly protein-coding (81.6%; **Figure 5B, Supplementary Figure 3E**), consistent with the ADAR2-shifted stoichiometry in the aggressive line.

**Figure 5.**
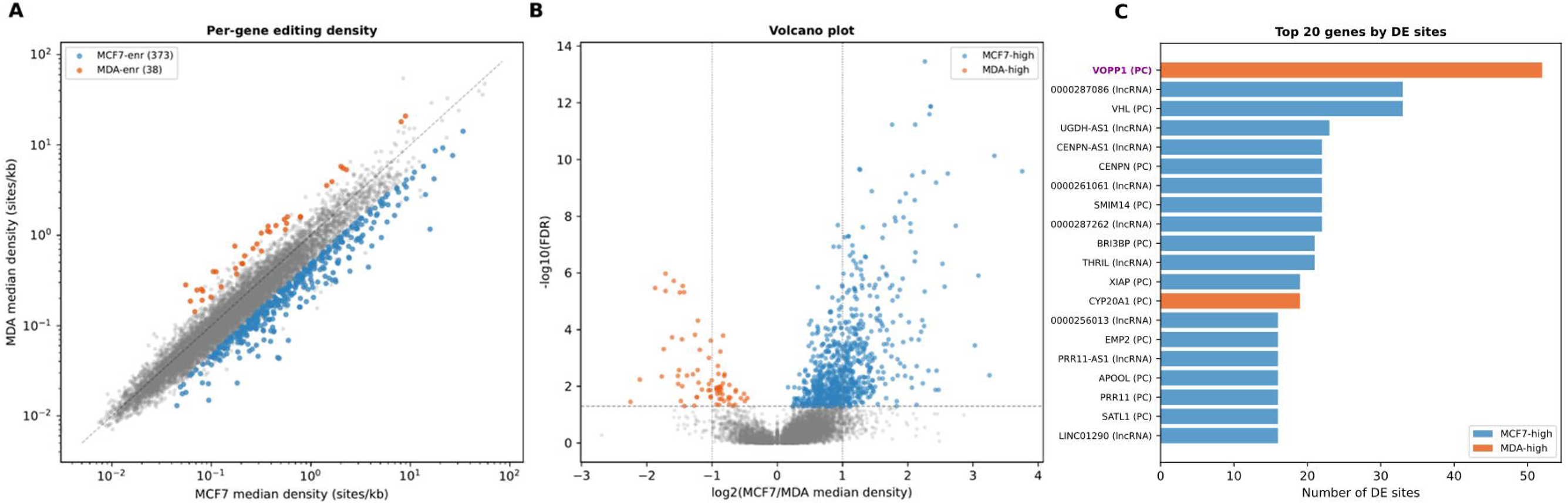
Subtype-specific gene editing programs and pathway enrichment **(A).** Subtype-specific gene editing programs. Log-log scatter of MCF7 versus MDA-MB-231 median per-sample editing density (sites/kb) for 9,538 protein-coding and lncRNA genes tested in both cell lines. Blue: MCF7-enriched genes (n = 373; FDR < 0.05, fold ≥ 2); orange: MDA-MB-231-enriched (n = 38); grey: non-significant. The strong asymmetry (373 vs 38 enriched genes) is consistent with MCF7’s higher ADAR1 expression driving broader editing. **(B)** Volcano plot showing log₂ fold change (MCF7/MDA-MB-231 median density) versus −log₁₀(FDR) for all tested genes. Dashed horizontal line: FDR = 0.05; dashed vertical lines: ±1 (2-fold). **(C)** Top 20 genes ranked by total DE sites, with MCF7-high (blue) and MDA-high (orange) sites shown as stacked bars. *VOPP1*, a known NF-κB activating oncogene at 7p11.2, harboured the most DE sites (52, all MDA-high).

#### Differentially edited sites cluster in gene-level programs

To identify individual editing sites with significantly different detection frequencies between cell lines, we performed differential editing analysis using DESeq2, treating per-site sample occurrence as a count variable with cell line as the design factor. This yielded 7,074 significant editing sites (padj < 0.05), which we mapped to genes via VEP annotation. Of these, 4,945 were MCF7-high (negative log2FC) and 2,129 MDA-high (positive log2FC). After mapping to genes, 931 genes harbored at least one differentially edited site (**Supplementary Figure 3G**), with 349 genes containing ≥3 sites. The direction consistency was remarkably high (mean 0.98): when a gene had differential editing, nearly all its sites shifted in the same direction, arguing against random noise (**Supplementary Figure 3F**). The gene with the most differential sites was *VOPP1* (52 sites, all MDA-high, mean log2FC = +2.59; **Figure 5C**), a known NF-κB activating oncogene located at 7p11.2, a region frequently amplified in aggressive breast cancers.

#### Fatty acid metabolism pathways are enriched among differentially edited antisense gene pairs

To investigate pathways associated with differential editing at sense-antisense loci, we performed over-representation analysis (ORA) on genes from antisense gene pairs with significantly different editing levels between cell lines. Among genes with higher editing in the MDA-MB-231 cell line, ORA revealed significant enrichment for fatty acid metabolism pathways, including alpha-linolenic (omega-3) and linoleic (omega-6) acid metabolism (Reactome R-HSA-2046104 and R-HSA-2046106; FDR = 0.003), omega-3/omega-6 fatty acid synthesis (WikiPathways; FDR = 0.004), and sebaleic acid formation (FDR = 0.008). These pathways converged on three key genes: FADS1, FADS2, and ACOX1, all of which participate in polyunsaturated fatty acid biosynthesis. No significant pathway enrichment was detected among genes with higher editing in MCF7.

The enrichment of lipid metabolism pathways among MDA-MB-231-enriched antisense editing targets is consistent with the established role of lipid metabolic reprogramming in triple-negative breast cancer. Lipid dysregulation, particularly in polyunsaturated fatty acid metabolism, has been shown to promote tumor growth and metastasis, and phosphatidylethanolamines have been identified as TNBC aggressiveness indicators by comparing MCF7 and MDA-MB-231 cell lines. The convergence of A-to-I editing differences at sense-antisense loci onto fatty acid metabolism pathways suggests a potential epitranscriptomic link between antisense-mediated editing regulation and the lipid metabolic phenotype characteristic of aggressive breast cancer.

### *NDUFS1* and *NDUFS1*-*AS1*: a candidate sense–antisense pair for experimental validation

#### In vitro validation of NDUFS1 editing

When integrating results across all computational analyses, *NDUFS1* (NADH:ubiquinone oxidoreductase core subunit S1) emerged as a compelling candidate based on several convergent lines of evidence. First, *NDUFS1* was constitutively and heavily edited in both cell lines, with a median of 96 editing sites per sample in MCF7 and 80 in MDA-MB-231, detected in 97.8% and 98.7% of samples respectively. Despite comparable overall editing density (2.15 vs 1.79 sites/kb; Mann-Whitney p = 0.33), *NDUFS1* ranked among the top 25 genes by number of differentially edited sites, harboring 14 positions with significantly higher editing in MCF7 (all MCF7-high; mean log2FC = −1.63, DESeq2 padj < 0.05). This pattern - comparable total editing load but divergent site-specific editing - is consistent with the model in which local dsRNA structure, potentially modulated by antisense transcription, determines which specific positions are edited rather than the overall editing capacity of the gene. Second, *NDUFS1* overlaps a novel lncRNA on the opposite strand (ENST00000453039), which we designated *NDUFS1*-*AS1*. This natural antisense transcript generates a sense–antisense overlap region where intermolecular dsRNA can form, providing a plausible structural basis for the site-specific editing differences observed between cell lines: differential expression of *NDUFS1*-*AS1* could alter the local dsRNA landscape at specific positions within *NDUFS1*, modulating ADAR access to individual editing sites without affecting the gene’s overall editing density (**Figure 6A; Supplementary Figure 4A-B)**. Third, *NDUFS1* encodes a core subunit of mitochondrial complex I, linking editing regulation to oxidative phosphorylation - a pathway known to be differentially regulated between ER+ and TNBC subtypes. These properties make the *NDUFS1*/*NDUFS1*-*AS1* pair an ideal model for experimental investigation of antisense-dependent editing regulation, which we pursued through *in vitro* validation described below.

**Figure 6.**
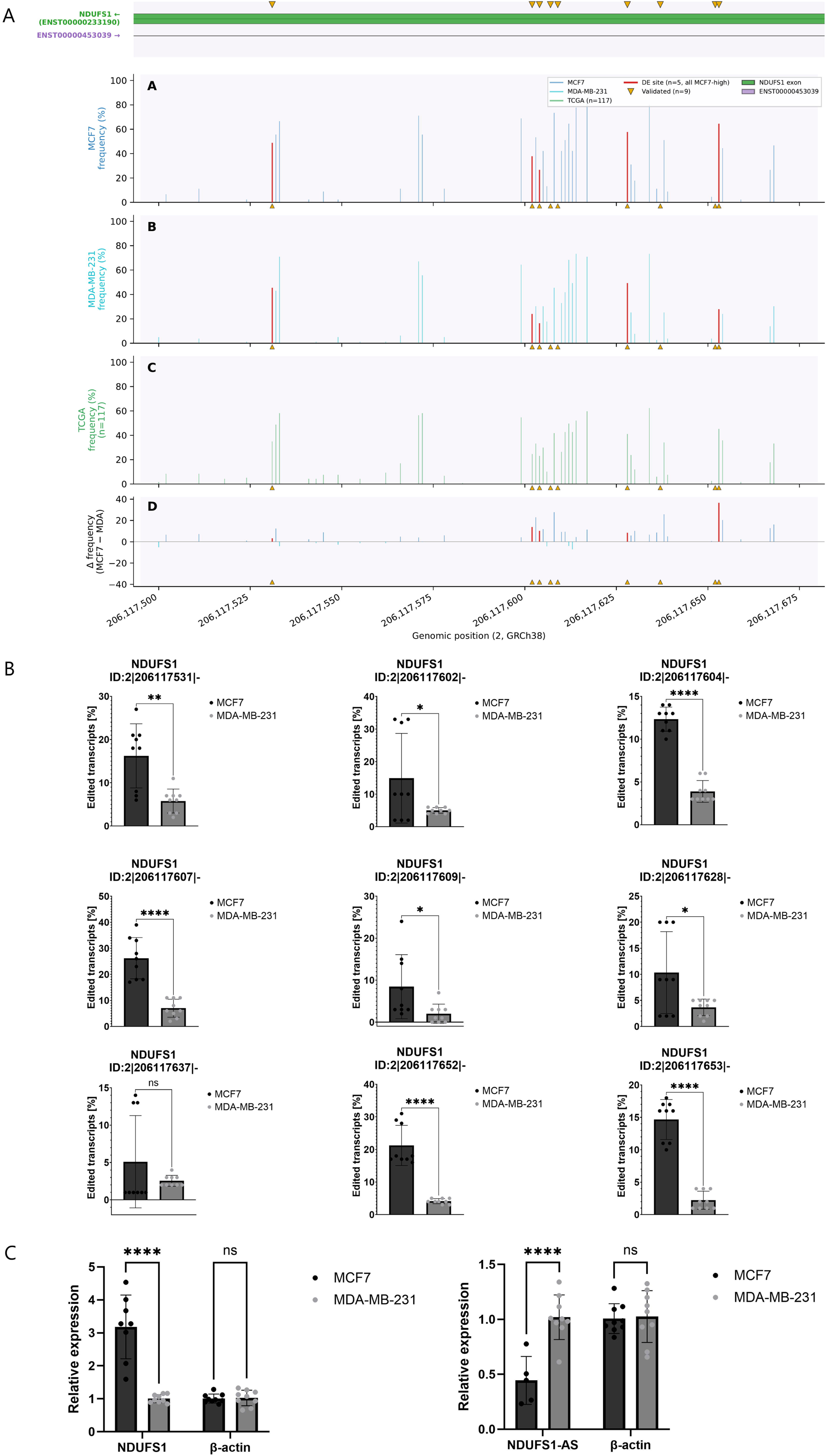
*NDUFS1* and *NDUFS1*-*AS1*: a candidate sense–antisense pair for experimental validation **(A)** Validated editing cluster at the *NDUFS1*/*NDUFS1*-*AS1* overlap boundary. High-resolution view of the editing cluster at 93.6–94.0% of the *NDUFS1* gene body (chr2:206,117,400–206,117,700), the region targeted for Sanger sequencing validation. Yellow arrowheads mark nine experimentally validated A-to-I editing positions (chr2:206,117,531–206,117,653). All validated sites overlap with computationally identified editing events detected in both cell lines, with MCF7 showing higher detection frequencies at the differentially edited positions. This cluster falls within the sense–antisense overlap region (purple shading), consistent with the model in which NDUFS1-AS1 transcription modulates local dsRNA structure and ADAR access at specific positions. **(B)** A-to-I RNA editing levels in two breast cancer subtypes. MCF7 cell line represents ER+ subtype and MDA-MB-231 TNBC subtype. Editing levels were calculated using EditR tool by analysing chromatograms from Sanger sequencing of PCR products. Data were presented as mean ± standard deviation (SD) from at least three independent experiments. Each plot shows one A-to-I RNA editing *locus* in *NDUFS1* gene. **(C)** Differences in relative gene expressions between two breast cancer cell lines. (a). RT-qPCR results showing relative expression of *NDUFS1* gene with *ACTB* control. (b). RT-qPCR results showing relative expression of *NDUFS1*-AS gene with *ACTB* control.

To validate expression of both sense and antisense transcripts, RT-PCR was performed in MCF7 and MDA-MB-231 cell lines. We detected robust expression of *NDUFS1* in both cell lines, as well as expression of *NDUFS1-AS1*, confirming co-expression of sense–antisense transcript pairs at the locus (**Supplementary Figure 4D**). To validate RNA editing at the *NDUFS1* locus, Sanger sequencing of RT-PCR products was performed. Analysis of chromatograms confirmed the presence of A-to-I editing at nine distinct sites within the *NDUFS1* transcript (**Supplementary Figure 4C**).

Quantification of editing levels using EditR (Kluesner et al., 2018) demonstrated reproducible editing across multiple sites within the *NDUFS1* transcript. Notably, editing levels measured by comparing mean base editing efficiency based on Sanger sequencing chromatograms differed between breast cancer subtypes (**Figure 6B**), with distinct editing patterns observed between MCF7 and MDA-MB-231 cells. These differences were consistent across biological replicates and suggest subtype-specific regulation of RNA editing at this locus, moreover for five of those sites our bioinformatic analysis had similar results.

#### Relative genes expression differences between breast cancer subtypes

To further characterise the relationship between RNA editing and transcript abundance, we quantified the expression levels of both *NDUFS1* and its antisense transcript *NDUFS1-AS1* using RT-qPCR. We observed a reciprocal, subtype-specific expression pattern between the two transcripts. *NDUFS1* was expressed at higher levels in MCF7 cells, whereas *NDUFS1-AS1* was upregulated in MDA-MB-231 cells (**Figure 6C**). This opposing expression pattern suggests differential regulation of sense and antisense transcription between breast cancer subtypes and has important implications for RNA editing. In MCF7 cells, higher *NDUFS1* expression in the context of elevated global ADAR1 activity is consistent with a dominant, intramolecular editing program driven by abundant Alu-derived dsRNA structures. In contrast, increased *NDUFS1-AS1* expression in MDA-MB-231 cells may facilitate the formation of intermolecular sense–antisense RNA duplexes, providing additional, locus-specific substrates for ADAR-mediated editing.

## Materials and Methods

### RNA-seq data acquisition

Publicly available RNA-seq datasets were retrieved from the NCBI Sequence Read Archive (SRA) (Leinonen et al., 2011) for two breast cancer cell lines: MCF7 (representing the ER+ luminal A subtype; 57 samples) and MDA-MB-231 (representing the triple-negative basal subtype; 101 samples). Of these, 34 samples were excluded due to a low number of detected A>I editing sites (fewer than 10,000). Sample accession numbers are listed in **Supplementary Table 1**. An additional 118 breast cancer patient tumour RNA-seq samples were obtained under controlled access from the database of Genotypes and Phenotypes (dbGaP; Breast Cancer Genome Guided Therapy Study (BEAUTY)) (Mailman et al., 2007). One sample (SRR3069831) was excluded based on the same criterion, with a higher threshold applied (fewer than 20,000 sites) to account for the generally greater number of A>I editing sites detected in patient-derived samples. All datasets consisted of paired-end Illumina RNA-seq reads.

### Read preprocessing and quality control

Raw sequencing reads were downloaded using the SRA Toolkit (fasterq-dump) and subjected to quality control and adapter trimming using fastp v0.23 (Chen et al., 2018) with default parameters. Reads shorter than 36 bp after trimming were discarded. Quality metrics were assessed using FastQC v0.12.1 (Andrews, 2010) and aggregated with MultiQC v1.14 (Ewels et al., 2016).

### Read alignment and gene expression quantification

Trimmed reads were aligned to the GRCh38 human reference genome (Ensembl release 110) (Martin et al., 2023) using STAR v2.7.10b (Dobin et al., 2013) in two-pass mode, with default parameters optimised for downstream editing site detection. The STAR genome index was generated using the corresponding Ensembl GTF annotation (Homo_sapiens.GRCh38.110.gtf). Alignment quality was verified by assessing the fraction of uniquely mapped reads; samples yielding fewer than 5 million uniquely mapped reads were excluded from downstream analysis. Gene-level expression was quantified using RSEM v1.3.3 (Li & Dewey, 2011) with the same genome and GTF annotation, producing per-sample transcripts per million (TPM) and expected count matrices. Library sizes (total reads mapped to genes) were extracted from the RSEM output for sequencing depth normalisation.

### Identification of A-to-I RNA editing sites

A-to-I RNA editing sites were identified using the SPRINT toolkit (Zhang et al., 2017), a reference-based, SNP-free approach that detects RNA editing events without requiring matched DNA sequencing data. SPRINT was run on each sample independently using the GRCh38 reference genome and RepeatMasker annotation of Alu elements (1,238,897 Alu elements) (Smit et al., 2013-2015). The SPRINT pipeline identifies candidate editing sites by detecting mismatches between RNA-seq reads and the reference genome, then applies a series of filters to distinguish genuine RNA editing from sequencing errors and genomic variants, including strand-specific mismatch filtering, multiple-testing correction, and the removal of sites in homopolymer regions. Only A-to-G mismatches on the sense strand and T-to-C mismatches on the antisense strand were retained as candidate A-to-I editing events. Per-sample editing site coordinates were output in BED format for downstream annotation.

### Variant annotation and filtering

Candidate editing sites were converted from BED to VCF format and annotated using the Ensembl Variant Effect Predictor (VEP, release 110) (McLaren et al., 2016) with the GRCh38 reference assembly. VEP was configured to report transcript-level consequences (e.g., intronic, UTR, missense, synonymous, splice site), existing variant identifiers (dbSNP, gnomAD, ClinVar), protein domain annotations, and allele frequencies from gnomAD exomes. VEP output fields included: Uploaded_variation, Location, Allele, Gene, Feature, Feature_type, Consequence, cDNA_position, CDS_position, Protein_position, Amino_acids, Codons, Existing_variation, DOMAINS, gnomADe_AF, CLIN_SIG, SOMATIC, and PHENO.

To distinguish genuine RNA editing from genomic variants, a stringent filtering pipeline was applied. Sites were removed if they: (i) were annotated in ClinVar (Landrum et al., 2020) with any clinical significance, (ii) had a gnomAD exome allele frequency ≥0.1% (Karczewski et al., 2020), or (iii) matched any dbSNP entry (Sherry et al., 2001). Only A>G and T>C substitutions consistent with A-to-I editing were retained in the final dataset (vep_filtered_permissive_v2). This filtering yielded 2,232,436 unique editing sites across all cell line samples.

### Alu element overlap analysis

The overlap between editing sites and Alu repetitive elements was computed using bedtools intersect v2.31.0 (Quinlan & Hall, 2010). Editing site coordinates were intersected with the RepeatMasker Alu annotation (GRCh38) in strand-agnostic mode. Per-cell-line and per-sample Alu fractions were computed as the number of editing sites overlapping Alu elements divided by the total number of editing sites. Statistical comparison of Alu fractions between cell lines was performed using Fisher’s exact test (for unique sites) and the Mann-Whitney U test (for per-sample distributions).

### ADAR enzyme expression analysis

Expression levels of ADAR family genes - ADAR (ADAR1; ENSG00000160710), ADARB1 (ADAR2; ENSG00000197381), and ADARB2 (ADAR3; ENSG00000185736) – were extracted from RSEM-derived TPM values for each sample. Between-cell-line comparisons were performed using two-sided Mann-Whitney U tests with Bonferroni correction. Effect sizes were quantified using Cohen’s d. The ADAR1-to-ADAR2 expression ratio was computed per sample and compared between cell lines. The relationship between ADAR1 expression and per-sample editing site count was assessed using Spearman rank correlation, both overall and within each cell line.

### Sequencing depth independence analysis

To verify that editing site counts were not confounded by sequencing depth, we computed Spearman correlations between library size (total reads mapped to genes from RSEM) and the number of unique editing sites per sample, both overall and within each cell line. Linear regression of editing site count on library size was performed across all samples, and residuals were compared between cohorts using the Kruskal-Wallis test. For the three-cohort analysis (MCF7, MDA-MB-231, patients), library sizes were compared using Mann-Whitney U tests and the combined regression was performed across all 240 samples.

### Functional distribution and sharing rate analysis

Each unique editing site was classified into a single functional category based on VEP consequence annotations using a priority hierarchy: splice site > high-impact coding (stop gained/lost, frameshift) > missense > synonymous > UTR > intronic > noncoding transcript > regulatory > upstream/downstream > intergenic. Sites classified as ‘intergenic’ by VEP represent editing events detected in RNA-seq reads mapping to positions outside annotated gene boundaries, likely reflecting unannotated transcripts or transcriptional read-through. When a site had multiple consequence annotations, the highest-priority category was assigned. The distribution of editing sites across functional categories was compared between cell lines using a chi-squared test of independence. Category-specific odds ratios and Fisher’s exact test p-values were computed for each functional category. The synonymous-to-missense ratio was compared between cell lines using Fisher’s exact test on the 2×2 contingency table of synonymous and missense site counts.

Variant sharing rates were computed per functional category as the mean percentage of samples within a cohort in which each variant was detected. Sharing rates were calculated independently for MCF7 (45 samples), MDA-MB-231 (79 samples), and patient (117 samples) cohorts. Overall sharing rates were computed as the variant-count-weighted mean across all categories.

### Core editome identification

Common (core) editing sites were defined as those present in ≥50% of samples within a cell line (≥23/45 for MCF7; ≥40/79 for MDA-MB-231). Cross-cell-line overlap of core editomes was quantified using two complementary approaches: (i) the fraction of core sites from one cell line detected in at least one sample of the other, and (ii) the number of sites independently classified as core in both cell lines. Expected overlap under the null hypothesis of random core status assignment was estimated analytically using the hypergeometric distribution and empirically via 10,000 random permutations of core labels across the 2,232,436 unique editing sites. A Jaccard index was computed as the size of the intersection divided by the size of the union of per-cell-line editing site sets.

### Sense–antisense overlap and editing density analysis

Sense–antisense overlapping regions were identified from the Ensembl GTF annotation (release 110) by intersecting gene coordinates on opposite strands using bedtools intersect (-S flag for opposite-strand). The non-redundant length of overlapping regions was computed by merging overlapping intervals per strand (bedtools merge) and summing interval lengths. Editing density was defined as the number of unique editing sites divided by the total nucleotide length of the region, expressed as a percentage.

Editing densities in overlapping versus non-overlapping regions were compared under four progressively stringent expression filters: (i) all annotated genes (no expression filter), (ii) actively transcribed genes (≥5 TPM in at least one sample), (iii) balanced expression ratio between sense and antisense partners (1:5 to 5:1), and (iv) tightly balanced expression (1:2 to 2:1). For each filter, a chi-squared test with Yates continuity correction was used to compare editing densities, and 95% Wilson confidence intervals were computed. The monotonic relationship between expression balance and editing enrichment was assessed using the Cochran-Armitage trend test.

### Antisense expression–editing correlation analysis

To test whether antisense transcript abundance quantitatively predicts editing efficiency, Spearman rank correlation coefficients were computed between the antisense:sense expression ratio and the editing ratio for each expressed gene pair. The editing ratio was defined as the number of editing sites multiplied by their editing levels, normalised by gene length. Only gene pairs where both partners had ≥10 TPM in at least 10 samples were included, yielding 332 testable lncRNA antisense sites and 1,947 protein-coding antisense sites.

The distribution of correlation coefficients was assessed for positive bias using the binomial test (proportion of positive ρ > 50%) and the one-sample t-test (mean ρ > 0). The proportion of positive correlations was compared between biotypes using Fisher’s exact test. Multiple testing correction was performed using the Benjamini–Hochberg procedure, and the number of significant results at FDR < 0.05 was compared to the expected number of false positives to assess enrichment.

### Gene-level antisense editing analyses

#### Approach A: Editing probability in genes with and without antisense lncRNA

Protein-coding genes were filtered for expression (≥1 TPM in at least one sample) and length (≥1 kb), yielding 12,653 genes. Genes were classified by the presence or absence of an overlapping antisense lncRNA based on the Ensembl GTF annotation. The proportion of edited genes (containing ≥1 editing site in ≥1 sample) was compared between groups using Fisher’s exact test, separately for each cell line. Among edited genes, per-sample editing density (sites per kilobase) was compared using the Mann-Whitney U test.

#### Approach B: Within-gene editing localization

For 1,411 protein-coding genes with partial antisense lncRNA overlap, editing density was compared between the overlap segment and position-matched control segments from 11,297 genes lacking antisense lncRNA. Position matching was performed by computing the fractional position of the overlap within the gene body (median: 10–55% from the 5′ end) and extracting the corresponding segment from control genes. Chi-squared tests were used to compare editing densities between overlap segments, position-matched controls, and non-overlap segments of the same genes.

#### Approach C: Overlap length dose–response

The relationship between sense–antisense overlap length and editing probability was assessed among 1,625 protein-coding genes with antisense lncRNA overlaps. Genes were binned by overlap length (200–500 bp, 500 bp–1 kb, 1–2 kb, 2–5 kb, 5–10 kb, 10–20 kb, 20–50 kb, >50 kb), and the fraction of edited genes was computed per bin. The Cochran–Armitage trend test was used to assess the monotonic relationship. Partial Spearman correlation between overlap length and editing probability, controlling for gene length, was computed using rank residuals. The overlap fraction (overlap length / gene length) was tested as an additional predictor.

#### Approach D: Alu element content in overlap regions

For each sense–antisense overlap, the number and orientation of Alu elements were determined by intersecting overlap coordinates with the RepeatMasker Alu annotation using bedtools. Overlaps were classified as: no Alu (neither strand), single-strand Alu (one strand only), or inverted Alu (both strands). Editing probability was compared across these categories using the Cochran–Armitage trend test and Fisher’s exact test (inverted Alu vs no Alu). The correlation between Alu element count and editing density was assessed using Spearman rank correlation.

#### Approach E: Intramolecular versus intermolecular dsRNA

To dissect the contributions of intramolecular dsRNA (inverted Alu pairs within a single transcript) and intermolecular dsRNA (formed between sense and antisense transcripts), all 14,195 expressed protein-coding genes were classified by two binary properties: presence/absence of inverted Alu pairs within the gene body, and presence/absence of an antisense lncRNA partner. A logistic regression model with an interaction term was fitted to editing status (edited vs not edited), and the interaction p-value was used to test whether the two mechanisms operate independently.

### Gene-level differential editing analysis

#### Per-sample editing density comparison

For each gene edited in ≥5 samples of both cell lines, per-sample editing density (sites per kilobase of gene length) was computed and compared between MCF7 and MDA-MB-231 using two-sided Mann-Whitney U tests. P-values were corrected for multiple testing using the Benjamini–Hochberg procedure (FDR < 0.05). Genes with ≥2-fold difference in median density were classified as MCF7-enriched or MDA-enriched. Biotype enrichment among differentially edited genes was assessed using Fisher’s exact test against the background of all tested genes.

#### Site-level differential editing with DESeq2

To identify individual editing sites with significantly different editing frequencies between cell lines, a count-based approach was implemented using DESeq2 v1.38 (Love et al., 2014). For each editing site, a per-sample count matrix was constructed where each entry represented the number of samples in which the site was detected (binary: 0 or 1). DESeq2 was run with cell line as the design variable, and sites with adjusted p-value < 0.05 were classified as differentially edited. Sites with negative log2 fold change were classified as MCF7-high, and those with positive log2FC as MDA-high. Differentially edited sites were mapped to genes via VEP annotation, and the number of DE sites per gene and direction consistency (fraction of sites shifting in the majority direction) were computed.

### Pathway enrichment analysis

Over-representation analysis (ORA) was performed on gene lists derived from the differential editing analysis using the ConsensusPathDB web server (CPDB; Herwig et al., 2016). Two separate analyses were conducted: (i) genes with higher overall editing levels in MCF7 versus MDA-MB-231, and (ii) genes with higher editing in MDA-MB-231 versus MCF7, using the full set of expressed genes as background. Pathway databases queried included Reactome, WikiPathways, and KEGG. P-values were corrected using the Benjamini–Hochberg method, and pathways with FDR < 0.01 were reported.

Additionally, targeted ORA was performed on genes from sense–antisense pairs with significantly different editing levels between cell lines. Up-regulated (MDA-high) and down-regulated (MCF7-high) gene lists were analysed separately using CPDB with Reactome and WikiPathways databases.

### NDUFS1 candidate gene selection criteria

*NDUFS1* was selected for experimental validation based on integration of results from all computational analyses. Selection criteria included: (i) constitutive editing in both cell lines (detected in >97% of samples), (ii) high editing density (>1.5 sites/kb), (iii) presence of differentially edited sites between cell lines (DESeq2 padj < 0.05), (iv) overlap with a natural antisense transcript (ENST00000453039, designated *NDUFS1*-*AS1*) providing a structural basis for dsRNA-dependent editing, and (v) biological relevance as a core subunit of mitochondrial complex I linking editing regulation to oxidative phosphorylation. Editing sites within the *NDUFS1* locus were visualised using a custom Python script that displays per-sample editing frequencies, exon/intron structure, antisense transcript overlap, Alu element positions, and experimentally validated sites.

### Genome annotation and repeat element resources

All analyses used the GRCh38/hg38 human reference genome assembly. Gene annotations were from Ensembl release 110 (Homo_sapiens.GRCh38.110.gtf). Repeat element annotations (Alu elements) were obtained from the UCSC Genome Browser RepeatMasker track. Sense–antisense overlapping regions were computed using the merged gene coordinate file (genes.merged.5.bed) derived from the Ensembl GTF by merging overlapping exonic and intronic regions per gene. Genome arithmetic operations (intersection, merge, complement, coverage) were performed using bedtools v2.31.0.

### Computational environment and statistical analysis

All bioinformatics analyses were performed on an Ubuntu 22.04 workstation. Python v3.10 (Python Software Foundation) was used for custom analysis scripts, with dependencies including NumPy v1.24 (Harris et al., 2020), pandas v2.0 (McKinney, 2010), SciPy v1.11 (Virtanen et al., 2020), statsmodels v0.14 (Seabold & Perktold, 2010), and Matplotlib v3.8 (Hunter, 2007). R v4.3 (R Core Team, 2023) was used for DESeq2 analysis. Genome arithmetic was performed with bedtools v2.31.0. Read alignment used STAR v2.7.10b and expression quantification used RSEM v1.3.3. Variant annotation used Ensembl VEP release 110. Pathway enrichment was performed using ConsensusPathDB (accessed March 2025).

All statistical tests were two-sided unless otherwise specified. Multiple testing correction was performed using the Benjamini–Hochberg procedure where indicated, with significance thresholds of FDR < 0.05 for genome-wide comparisons and nominal p < 0.05 for single-gene or targeted analyses. Effect sizes are reported as odds ratios (for binary outcomes), Spearman ρ (for correlations), or Cohen’s d (for between-group comparisons). 95% confidence intervals are Wilson intervals (for proportions) or bootstrap intervals (for medians). Publication-quality figures were generated using Matplotlib with a consistent colour palette: MCF7 = #3182bd (steel blue), MDA-MB-231 = #17becf (teal), patient = #31a354 (green), DE sites = #d62728 (red).

### Cell culture

MCF7 and MDA-MB-231 human breast cancer cell lines were purchased by an ATCC supplier (ATCC, Manassas, VA, USA). Cell lines were maintained in RPMI medium (Thermo Scientific™, Waltham, MA, USA), supplemented with 10% fetal bovine serum (FBS) (Capricorn Scientific, Ebsdorfergrund, Germany) and Antibiotic-Antimycotic (100X) solution (Thermo Scientific™, Waltham, MA, USA) to prevent microbial contamination. Cell cultures were incubated in 37°C and an atmosphere of 5% CO2 in a humified incubator to ensure optimal conditions.

### RNA and DNA extraction

Total RNA and genomic DNA were extracted from the cells with TRI Reagent (Molecular Research Center, Cincinnati, OH, USA) according to the manufacturer’s protocol. The concentration and quality of isolated acids were checked using Spectrophotometer (DeNovix, Wilmington, DE, USA) by measuring absorbance ratios at 260/280 nm and 260/230 nm.

Additionally, RNA’s parameters were improved by using RNA Clean & Concentrator™ (ZYMO Research, Irvine, CA, USA) kit with an additional DNase treatment step to eliminate potential DNA contamination. Purified RNA was stored at −80°C.

### Reverse transcription and PCR/qPCR

RNA was reverse transcribed with RevertAid First Strand cDNA Synthesis Kit (Thermo Scientific™, Waltham, MA, USA) according to the manufacturer’s instructions with optional step of heating the RNA with random hexamers for five minutes in 65°C to disrupt secondary structures. Synthesised cDNA was used for both PCR and quantitative PCR (qPCR) reactions. Primers for PCR reactions were designed to amplify specific regions of A-to-I RNA editing and qPCR primers were designed to not increase 200nt long products. Both types were designed using PrimerBlast tool. PCR reactions were performed with Phusion High Fidelity Polymerase (Thermo Scientific™, Waltham, MA, USA) in order to minimise the possibility of polymerase errors, which is especially important for studying single nucleotide changes within transcripts. qPCR reactions were done using SYBR Green PCR Master Mix (Thermo Scientific™, Waltham, MA, USA) and relative gene expression was calculated using the ΔΔCt method, normalizing to β-actin housekeeping gene. Absolute copiers of transcripts in both cell lines were measured based on standard qPCR curve in QuantStudio™ Real-Time PCR Software.

### Sanger sequencing and A-to-I RNA editing quantification

Products of PCR reactions were cleaned with DNA Clean-up and Concentration™ (ZYMO Research, Irvine, CA, USA) kit. Sanger sequencing was performed with reverse primers to improve precision and accuracy of detecting A-to-I RNA editing events (Eggington et al., 2011; Rinkevich et al., 2012). Measurements of A-to-I RNA editing levels were performed using EditR software (version 1.0.10) (Kluesner et al., 2018). EditR is an online tool that can be used to detect and quantify base editing from Sanger sequencing chromatograms.

### Statistical analysis

Statistical analyses were performed using GraphPad Prism (version 10.4.1). Statistical differences were calculated with ANOVA and t-test, p < 0.05 was considered as statistically significant. All data presents the mean ± standard deviation (SD) from at least three replicates.

## Discussion

A-to-I RNA editing, catalysed principally by ADAR1, is one of the most pervasive post-transcriptional modifications in the human transcriptome, with millions of sites concentrated in Alu repeat elements (Nishikura, 2016; Eisenberg & Levanon, 2018). While intramolecular dsRNA structures formed by inverted Alu pairs have been the primary focus of mechanistic studies, the contribution of intermolecular dsRNA arising from natural antisense transcription has received little attention. Here we demonstrate that ADAR1 expression is the dominant driver of subtype-divergent editing landscapes, that a conserved constitutive editome of ∼2,500 sites is maintained across subtypes, and that antisense lncRNA transcription constitutes an independent, additive pathway governing editing probability through intermolecular dsRNA formation. Experimental validation at the *NDUFS1*/*NDUFS1*-*AS1* locus provides cellular evidence supporting this model.

### ADAR1 expression as the primary determinant of subtype-specific editing breadth

The higher editing site count in MCF7 is explained in large part by its elevated ADAR1 expression, consistent with evidence that ADAR1 is the dominant editor of Alu-derived dsRNA and that its overexpression predicts editing output across cancer types (Chen et al., 2013; Han et al., 2015; Paz-Yaacov et al., 2015; Fumagalli et al., 2015). The higher ADAR1/ADAR2 ratio in MCF7 likely drives a predominantly intronic Alu-targeting programme, whereas the relatively greater ADAR2 contribution in MDA-MB-231 shifts editing toward synonymous positions - potentially influencing translation kinetics, as inosine at synonymous positions can slow ribosome elongation (Licht et al., 2019) - without altering protein sequence. ADAR2 is well-known for highly site-selective editing of coding sequences at positions not productively targeted by ADAR1 (Nishikura, 2016).

The absence of a within-cell-line ADAR1–editing correlation in MCF7, contrasting with the modest positive correlation in MDA-MB-231, is consistent with a limiting dsRNA substrate pool: once ADAR1 exceeds the level required to edit all accessible targets, further increases do not expand the editome (Walkley & Li, 2017; Liddicoat et al., 2015).

### A constitutive core editome persists across divergent editing landscapes

The 230-fold enrichment of shared core sites over chance implies that these loci possess unusually high-affinity ADAR substrates, consistent with evidence that near-perfect Alu-pair dsRNA hairpins are processed with substantially more favourable kinetics than imperfect structures (Lehmann & Bass, 1999). The concentration of patient core sites in UTRs (55.5%) is notable given that ADAR1 binding to 3′ UTRs can stabilise mRNAs by displacing destabilising factors (Wang et al., 2013; Bahn et al., 2015), as also reflected in RADAR and REDIportal databases (Ramaswami & Li, 2014; Picardi et al., 2017). The functional sharing hierarchy (UTR > splice site > noncoding transcript > regulatory > intronic > intergenic) was preserved across all three cohorts despite their markedly different ADAR1 expression levels, suggesting that the reproducibility of editing at a given site is determined by local transcript structure - such as dsRNA stability and ADAR accessibility - rather than by the overall abundance of the editing enzyme.

### Antisense lncRNA transcription as an independent editing pathway through intermolecular dsRNA

A previous study by Kawahara and Nishikura examined sense–antisense overlapping regions genome-wide and found that editing was depleted rather than enriched at these loci, leading them to conclude that intermolecular dsRNA formation between sense and antisense transcripts is rare in vivo (Kawahara & Nishikura, 2006). Our data replicate this depletion (0.66-fold) when all annotated overlapping regions are analysed regardless of expression. However, the mere presence of two genes on opposite strands does not guarantee that both are actively transcribed in the same cell - and dsRNA can only form if both transcripts are present simultaneously. When we restricted the analysis to overlapping gene pairs where both partners are actively co-expressed at balanced levels, the pattern reversed to a 1.55-fold enrichment of editing. This indicates that the earlier null result was driven by the inclusion of silent or lowly expressed gene pairs that produce no intermolecular dsRNA despite their genomic co-location.

The positive directional bias among both lncRNA and protein-coding antisense sites, with no biotype difference, supports a purely structural model in which any antisense RNA can form intermolecular dsRNA and present an ADAR substrate. The gene-level analyses provide a mechanistic framework: antisense overlap governed editing probability but not density, consistent with evidence that dsRNA access - rather than catalytic capacity - determines editing outcomes (Quinones-Valdez et al., 2019). The within-gene comparison showing higher editing density in overlap segments controls for all gene-level confounders. The dose–response with overlap length is consistent with promiscuous ADAR deamination of long dsRNA substrates (Polson & Bass, 1994), and the steep Alu gradient within overlaps reflects that Alu sequences in both transcripts enable complementary intermolecular base-pairing, amplifying the intramolecular substrates characterised in earlier studies (Bazak et al., 2014b).

The 2×2 dissection establishes a clear hierarchy: inverted Alu pairs are the dominant determinant (OR = 36.5) while antisense lncRNA provides an independent, additive contribution (OR ≈ 1.5) with no interaction. To our knowledge, this is the first genome-wide demonstration that antisense transcription scales editing probability in proportion to overlap length and Alu content, extending isolated examples (Pecori et al., 2022) to a broadly operative mechanism.

Caveats include the possibility that ADAR editing stabilises antisense transcripts rather than the reverse, though the within-gene spatial restriction, Alu requirements, and *NDUFS1* data make this less parsimonious. Other regulatory inputs - including RBP competition (Quinones-Valdez et al., 2019) and ADAR post-translational modifications such as phosphorylation and SUMOylation (Sakurai et al., 2017; Desterro et al., 2003; Desterro et al., 2005) - likely further modulate editing at individual loci.

### Experimental validation at the *NDUFS1*/*NDUFS1*-*AS1* locus

*NDUFS1* was selected for validation based on constitutive editing in both cell lines, the presence of differentially edited positions, and the structural overlap with a novel antisense transcript (ENST00000453039, designated *NDUFS1*-*AS1*). RT-PCR confirmed co-expression of *NDUFS1*, *NDUFS1*-*AS1*, and the *ACTB* control in both MCF7 and MDA-MB-231, establishing the cellular prerequisite for intermolecular dsRNA formation. *NDUFS1*-*AS1* has not previously been characterised; its consistent amplification across independent experiments confirms it is not a cell-culture artefact.

Sanger sequencing of RT-PCR products using reverse primers - a strategy validated for more accurate A-to-I editing detection (Rinkevich et al., 2012) - revealed T-to-C changes (reverse complement of A-to-G) at multiple cDNA positions within the 3′-terminal region of *NDUFS1*. Matched genomic DNA from both cell lines showed no corresponding differences, confirming post-transcriptional RNA editing rather than DNA variation - a critical orthogonal control for editome analyses (Eisenberg & Levanon, 2018). EditR quantification (Kluesner et al., 2018) of editing levels at individual loci revealed differential editing between the two cell lines, with MCF7 exhibiting higher editing at the validated positions, concordant with the computationally identified MCF7-high sites. Importantly, total *NDUFS1* editing density was comparable between cell lines while site-level editing differed - precisely the dissociation between density and site selectivity predicted by our antisense-modulated dsRNA model.

RT-qPCR demonstrated differential expression of both *NDUFS1* and *NDUFS1*-*AS1* between cell lines, consistent with the prediction that differences in antisense lncRNA abundance should accompany differences in site-specific editing. *NDUFS1* encodes a core subunit of mitochondrial complex I, and oxidative phosphorylation is differentially regulated between ER+ and TNBC subtypes (Sotgia et al., 2012), suggesting the editing differences may reflect or contribute to this metabolic divergence through ADAR-mediated modulation of transcript stability (Wang et al., 2013).

### Subtype-specific editing programs and oncogenic connections

Differential editing showed a striking gene-level asymmetry consistent with ADAR1-driven broad editing in MCF7. The most-densely edited genes were predominantly lncRNAs, consistent with their higher Alu repeat density due to weaker purifying selection (Bazak et al., 2014b). *VOPP1* harboured the most differentially edited sites (52, all MDA-MB-231-high), with extreme direction consistency. This oncogene suppresses apoptosis via WWOX sequestration (Bonin et al., 2018), activates NF-κB (Park & James, 2005), and is co-amplified with *EGFR* at 7p11.2. *VOPP1* joins oncogene transcripts undergoing cell-type-specific editing, including *AZIN1* in hepatocellular carcinoma (Chen et al., 2013) and GLI1 in multiple myeloma (Lazzari et al., 2017).

Pathway enrichment identified fatty acid metabolism among MDA-MB-231-enriched antisense editing targets, converging on FADS1, FADS2, and ACOX1 - genes whose elevated expression marks TNBC subsets with greater metastatic potential and ferroptosis vulnerability (Lorito et al., 2024). This raises the possibility that antisense-mediated epitranscriptomic regulation contributes to the lipid metabolic phenotype of aggressive breast cancer.

### Broader implications and limitations

This model has several implications. First, the editome is shaped not only by ADAR expression and intramolecular structure but also by the lncRNA landscape; given the tissue-specificity of lncRNA profiles (Derrien et al., 2012), this could contribute to cell-type specificity of editing across normal tissues (Tan et al., 2017). Second, intermolecular dsRNA may be the primary editing substrate at genes lacking inverted Alu pairs, explaining ADAR editing at such loci (Levanon et al., 2004; Kawahara & Nishikura, 2006). Third, lncRNA dysregulation in breast cancer (Gutschner & Diederichs, 2012) may have downstream epitranscriptomic consequences not previously considered.

The analyses rely on RNA-seq without matched genomic DNA, although the SPRINT toolkit was specifically designed to identify RNA editing sites from RNA-seq data alone by using intrinsic strand-specific and statistical filters to distinguish editing from genomic variants. Nevertheless, the 2.2 million sites should be interpreted as a candidate editome validated at nine *NDUFS1* positions by Sanger sequencing. Patient tumour analysis is limited by batch effects, and the cell lines represent extreme phenotypic poles. Causal proof that *NDUFS1*-*AS1* drives editing differences requires ASO-mediated knockdown.

## Conclusions

A-to-I RNA editing in breast cancer is shaped by a two-tier architecture: a dominant ADAR1-driven programme targeting intramolecular Alu dsRNA, upon which an independent lncRNA antisense pathway is superimposed via intermolecular dsRNA. The result is a subtype-specific editome preserving a constitutive core but diverging in magnitude, functional distribution, and site selection. Experimental validation at the *NDUFS1*/*NDUFS1*-*AS1* locus - including co-expression of sense and antisense transcripts, Sanger-confirmed editing at computationally predicted positions, and differential editing between MCF7 and MDA-MB-231 - supports the computational model. The convergence of differential antisense-locus editing onto fatty acid metabolism and oncogenic survival pathways points to an underappreciated epitranscriptomic dimension of breast cancer subtype biology.

## Acknowledgements

The computations were partially conducted at the Poznan Supercomputing and Networking Center.

## Author contributions

Conceptualization, M.W.S.; methodology, E.W. and M.W.S.; formal analysis, M.W.S. and K.S.; investigation, K.S. and E.W.; data curation, K.S. and M.W.S.; visualization, M.W.S and K.S.; supervision, E.W. and M.W.S.; funding acquisition, K.S., E.W and M.W.S. All authors participated in the writing, review, and editing of the manuscript. All authors read and approved the final version of the manuscript.

## Supplementary Materials

**Supplementary Figure 1.** Patient tumor editomes cluster distinctly from cell lines but preserve the functional sharing hierarchy. Mean sharing rates by functional category across all three cohorts. Sharing rate is defined as the percentage of samples within a cohort in which a given variant is detected, averaged across all variants in each category. Categories are ordered by MCF7 sharing rate (descending). The functional hierarchy - UTR (9.4%, 6.6%, 6.1%) > splice site (6.1%, 4.0%, 4.1%) > noncoding transcript > regulatory > intronic > intergenic - is preserved across all three cohorts, with MCF7 consistently showing the highest sharing rates, followed by MDA-MB-231 and patients. This graded decrease is consistent with greater inter-tumor heterogeneity diluting per-variant sharing in patients, while the rank order demonstrates that functionally constrained editing sites are shared more reproducibly than intronic or intergenic sites across breast cancer contexts.

**Supplementary Figure 2. Antisense expression correlates positively with editing efficiency independent of transcript biotype.** Top row: lncRNA antisense partners; bottom row: protein-coding antisense partners. **(Left)** Distribution of Spearman correlation coefficients (ρ) between antisense:sense expression ratio and editing ratio across all tested sites. Among 332 lncRNA sites, 182 (54.8%) exhibited positive correlations (binomial p = 0.019; mean ρ = 0.098); among 1,947 protein-coding sites, 1,053 (54.1%) were positive (binomial p = 2.2 × 10⁻⁵; mean ρ = 0.059). Dashed lines indicate mean (red) and median (orange) ρ values. **(Center)** Nominal p-value distributions. The strong enrichment of low p-values above the uniform null expectation (red dashed line) confirms genuine signal beyond what would be expected by chance. **(Right)** Distribution of Spearman ρ for sites surviving Benjamini–Hochberg correction at FDR < 0.10. The positive bias strengthened after multiple testing correction: 66.7% of lncRNA correlations (42/63; mean ρ = 0.337) and 64.5% of protein-coding correlations (218/338; mean ρ = 0.191) were positive.

**Supplementary Figure 3. (A)** Editing localises to the region of sense–antisense interaction. Distribution of antisense overlap start positions relative to gene body (0 = 5′, 1 = 3′). **(B)** Editing probability scales with antisense overlap length. Editing density within the overlap region versus overlap length among edited genes. **(C)** Total gene editing density versus overlap fraction of gene length; red points indicate binned medians. **(D)** Inverted Alu elements mediate editing at sense–antisense overlaps. Editing density as a function of Alu element count within the overlap region. **(E)** Subtype-specific gene editing programs. Biotype composition of the 411 differentially edited genes (373 MCF7-enriched + 38 MDA-enriched). MCF7-enriched genes were over-represented for lncRNAs (27.9% vs 17.0% background), while MDA-enriched genes were predominantly protein-coding (81.6%). **(F)** Differentially edited sites cluster in gene-level programs. MCF7-high versus MDA-high differentially edited sites per gene (≥3 DE sites; n = 349 genes). Points below the diagonal: MCF7-predominant (n = 266, blue); above: MDA-predominant (n = 81, orange). Direction consistency was 0.98, indicating coordinated gene-level shifts. **(G)** Distribution of DE sites per gene among 931 genes harbouring at least one differentially edited site. Dashed line: ≥3 threshold. **(H)** Editing landscape at the *VOPP1* locus. Per-sample editing site frequency across the 3′ region (80–90% of gene body) of *VOPP1* (ENST00000417399, chr7:12,632,291–12,695,890, minus strand) in MCF7 (A, blue), MDA-MB-231 (B, teal), and patient tumours (C, green). *VOPP1* harboured the most differentially edited sites of any gene in the dataset (52 sites, all MDA-MB-231-high, mean log₂FC = +2.59). The frequency difference panel (D) shows uniformly negative values, reflecting consistently higher editing in MDA-MB-231.

**Supplementary Figure 4. (A)** Editing landscape across the full *NDUFS1* locus. Per-sample editing site frequency across the *NDUFS1* gene (ENST00000233190, chr2:205,852,191–206,120,380, minus strand) in MCF7 (A, blue), MDA-MB-231 (B, teal), and patient tumours (C, green, n=117). Each vertical line represents one editing site; height indicates the percentage of samples in which the site was detected. Red lines denote differentially edited sites (DESeq2 padj < 0.05). The bottom panel (D) shows the difference in detection frequency between MCF7 and MDA-MB-231 (positive = MCF7-higher). The gene model (top) shows exon structure of the canonical transcript with the antisense lncRNA *NDUFS1*-*AS1* (ENST00000453039, purple) on the opposite strand; the purple shading marks the sense–antisense overlap region. Despite comparable overall editing density between cell lines (2.15 vs 1.79 sites/kb; p = 0.33), site-specific differences are concentrated in the 3′ region coinciding with the antisense overlap. **(B)** Editing landscape in the 3′ region of *NDUFS1* (80–100% of gene body). Zoomed view of the 3′-terminal 20% of the *NDUFS1* locus, encompassing the sense–antisense overlap with *NDUFS1*-*AS1*. This region harbours the majority of differentially edited sites, with editing frequencies consistently higher in MCF7 than MDA-MB-231 at DE positions (all MCF7-high; mean log₂FC = −1.63). Patient tumour editing frequencies at these positions were intermediate between the two cell lines. **(C)** Fragments of chromatograms of the studied *NDUFS1* gene with marked A-to-I RNA editing *loci*. To simplify the graph DNA from only one (of two) cell lines was shown. In both DNAs there were no significant differences. Exact *loci* are in black dotted boxes, red peaks (T) indicate no editing and blue one **(C)** that there is A-to-I change. The reason why chromatograms show T-to-C changes instead of A-to-G is the use of reverse primers in Sanger sequencing to increase the precision and accuracy of detecting RNA editing. **(D)** RT-PCR results performed on two breast cancer cell lines. In this experiment we have checked the expression of transcripts that undergo the process of A-to-I RNA editing, namely protein coding *NDUFS1* as well as its novel natural antisense transcript *NDUFS1*-*AS* (ENST00000453039) and *ACTB* control gene in MCF7 cell line (representing ER+ breast cancer subtype) and MDA-MB-231 (representing TNBC).

**Supplementary Table 1.** Cell line assignments for all MCF7 and MDA-MB-231 SRA samples considered in this study. Samples rejected due to low editing site counts (indicative of low sequencing quality or coverage) are indicated.

## References

1) Andrews S. FastQC: a quality control tool for high throughput sequence data. 2010.

2) Leinonen, R., Sugawara, H., Shumway, M., & on behalf of the International Nucleotide Sequence Database Collaboration. The Sequence Read Archive. Nucleic Acids Res 39, D19–D21 (2011).

3) Bahn, J. H. et al. Genomic analysis of ADAR1 binding and its involvement in multiple RNA processing pathways. Nat Commun 6, 6355 (2015).

4) Bazak, L. et al. A-to-I RNA editing occurs at over a hundred million genomic sites, located in a majority of human genes. Genome Res 24, 365–376 (2014a).

5) Bazak, L., Levanon, E. Y. & Eisenberg, E. Genome-wide analysis of Alu editability. Nucleic Acids Res 42, 6876–6884 (2014b).

6) Bonin, F. et al. VOPP1 promotes breast tumorigenesis by interacting with the tumor suppressor WWOX. BMC Biol 16, 109 (2018).

7) Brachova, P. et al. Inosine RNA modifications are enriched at the codon wobble position in mouse oocytes and eggs†. Biol Reprod 101, 938–949 (2019).

8) Chen, L. et al. Recoding RNA editing of AZIN1 predisposes to hepatocellular carcinoma. Nat Med 19, 209–216 (2013).

9) Chen, S., Zhou, Y., Chen, Y. & Gu, J. fastp: an ultra-fast all-in-one FASTQ preprocessor. Bioinformatics 34, i884–i890 (2018).

10) Delli Ponti, R., et al. A high-throughput approach to predict A-to-I effects on RNA structure indicates a change of double-stranded content in noncoding RNAs. IUBMB Life 75, 411–426 (2023).

11) Derrien, T. et al. The GENCODE v7 catalog of human long noncoding RNAs: analysis of their gene structure, evolution, and expression. Genome Res 22, 1775–1789 (2012).

12) Desterro, J. M. P. et al. Dynamic association of RNA-editing enzymes with the nucleolus. J Cell Sci 116, 1805–1818 (2003).

13) Desterro, J. M. P. et al. SUMO-1 modification alters ADAR1 editing activity. Mol Biol Cell 16, 5115–5126 (2005).

14) Dobin, A. et al. STAR: ultrafast universal RNA-seq aligner. Bioinformatics 29, 15–21 (2013).

15) Eggington, J. M., Greene, T. & Bass, B. L. Predicting sites of ADAR editing in double-stranded RNA. Nat Commun 2, 319 (2011).

16) Eisenberg, E. & Levanon, E. Y. A-to-I RNA editing - immune protector and transcriptome diversifier. Nat Rev Genet 19, 473–490 (2018).

17) Ewels, P., Magnusson, M., Lundin, S. & Käller, M. MultiQC: summarize analysis results for multiple tools and samples in a single report. Bioinformatics 32, 3047–3048 (2016).

18) Fumagalli, D. et al. Principles Governing A-to-I RNA Editing in the Breast Cancer Transcriptome. Cell Rep 13, 277–289 (2015).

19) Gutschner, T. & Diederichs, S. The hallmarks of cancer: a long non-coding RNA point of view. RNA Biol 9, 703–719 (2012).

20) Han, L. et al. The Genomic Landscape and Clinical Relevance of A-to-I RNA Editing in Human Cancers. Cancer Cell 28, 515–528 (2015).

21) Harris, C. R. et al. Array programming with NumPy. Nature 585, 357–362 (2020).

22) Herwig, R., Hardt, C., Lienhard, M. & Kamburov, A. Analyzing and interpreting genome data at the network level with ConsensusPathDB. Nat Protoc 11, 1889–1907 (2016).

23) Huang, W. et al. The snoRNA-like lncRNA LNC-SNO49AB drives leukemia by activating the RNA-editing enzyme ADAR1. Cell Discov 8, 117 (2022).

24) Hunter, J. Matplotlib: A 2D Graphics Environment. Computing in Science & Engineering 9, 90–95 (2007).

25) Karczewski KJ, Francioli LC, Tiao G, Cummings BB, Alföldi J, Wang Q, et al. The mutational constraint spectrum quantified from variation in 141,456 humans. Nature. 2020;581(7809):434–443.

26) Kawahara, Y. & Nishikura, K. Extensive adenosine-to-inosine editing detected in Alu repeats of antisense RNAs reveals scarcity of sense-antisense duplex formation. FEBS Lett 580, 2301–2305 (2006).

27) Kluesner, M. G. et al. EditR: A Method to Quantify Base Editing from Sanger Sequencing. CRISPR J 1, 239–250 (2018).

28) Landrum, M. J. et al. ClinVar: improvements to accessing data. Nucleic Acids Res 48, D835–D844 (2020).

29) Lazzari, E. et al. Alu-dependent RNA editing of GLI1 promotes malignant regeneration in multiple myeloma. Nat Commun 8, 1922 (2017).

30) Lehmann, K. A. & Bass, B. L. The importance of internal loops within RNA substrates of ADAR1. J Mol Biol 291, 1–13 (1999).

31) Levanon, E. Y. et al. Systematic identification of abundant A-to-I editing sites in the human transcriptome. Nat Biotechnol 22, 1001–1005 (2004).

32) Li, B. & Dewey, C. N. RSEM: accurate transcript quantification from RNA-Seq data with or without a reference genome. BMC Bioinformatics 12, 323 (2011).

33) Licht, K. et al. Inosine induces context-dependent recoding and translational stalling. Nucleic Acids Res 47, 3–14 (2019).

34) Liddicoat, B. J. et al. RNA editing by ADAR1 prevents MDA5 sensing of endogenous dsRNA as nonself. Science 349, 1115–1120 (2015).

35) Lorito, N. et al. FADS1/2 control lipid metabolism and ferroptosis susceptibility in triple-negative breast cancer. EMBO Mol Med 16, 1533–1559 (2024).

36) Love, M. I., Huber, W. & Anders, S. Moderated estimation of fold change and dispersion for RNA-seq data with DESeq2. Genome Biol 15, 550 (2014).

37) Mailman, M. D. et al. The NCBI dbGaP database of genotypes and phenotypes. Nat Genet 39, 1181–1186 (2007).

38) Martin, F. J., et al. Ensembl 2023. Nucleic Acids Res 51, D933–D941 (2023).

39) McKinney, W. Data Structures for Statistical Computing in Python. SciPy 2010 10.25080/Majora-92bf1922-00a (2010) doi:10.25080/Majora-92bf1922-00a.

40) McLaren, W. et al. The Ensembl Variant Effect Predictor. Genome Biol 17, 122 (2016).

41) Nishikura, K. A-to-I editing of coding and non-coding RNAs by ADARs. Nat Rev Mol Cell Biol 17, 83–96 (2016).

42) Park, S. & James, C. D. ECop (EGFR-coamplified and overexpressed protein), a novel protein, regulates NF-kappaB transcriptional activity and associated apoptotic response in an IkappaBalpha-dependent manner. Oncogene 24, 2495–2502 (2005).

43) Paz-Yaacov, N. et al. Elevated RNA Editing Activity Is a Major Contributor to Transcriptomic Diversity in Tumors. Cell Rep 13, 267–276 (2015).

44) Pecori, R. et al. ADAR RNA editing on antisense RNAs results in apparent U-to-C base changes on overlapping sense transcripts. Front Cell Dev Biol 10, 1080626 (2022).

45) Peng, X. et al. A-to-I RNA Editing Contributes to Proteomic Diversity in Cancer. Cancer Cell 33, 817–828.e7 (2018).

46) Picardi, E., D’Erchia, A. M., Lo Giudice, C. & Pesole, G. REDIportal: a comprehensive database of A-to-I RNA editing events in humans. Nucleic Acids Res 45, D750–D757 (2017).

47) Polson, A. G. & Bass, B. L. Preferential selection of adenosines for modification by double-stranded RNA adenosine deaminase. EMBO J 13, 5701–5711 (1994).

48) Quinlan, A. R. & Hall, I. M. BEDTools: a flexible suite of utilities for comparing genomic features. Bioinformatics 26, 841–842 (2010).

49) Quinones-Valdez, G. et al. Regulation of RNA editing by RNA-binding proteins in human cells. Commun Biol 2, 19 (2019).

50) R Core Team. R: A language and environment for statistical computing. R Foundation for Statistical Computing, Vienna, Austria. 2023.

51) Ramaswami, G. & Li, J. B. RADAR: a rigorously annotated database of A-to-I RNA editing. Nucleic Acids Res 42, D109–113 (2014).

52) Rinkevich, F. D., Schweitzer, P. A. & Scott, J. G. Antisense sequencing improves the accuracy and precision of A-to-I editing measurements using the peak height ratio method. BMC Res Notes 5, 63 (2012).

53) Sakurai, M. et al. ADAR1 controls apoptosis of stressed cells by inhibiting Staufen1-mediated mRNA decay. Nat Struct Mol Biol 24, 534–543 (2017).

54) Salameh, A. et al. PRUNE2 is a human prostate cancer suppressor regulated by the intronic long noncoding RNA PCA3. Proc Natl Acad Sci U S A 112, 8403–8408 (2015).

55) Samorowska, K., Wanowska, E. & Szcześniak, M. W. Novel lncRNA UGGT1-AS1 Regulates UGGT1 Expression in Breast Cancer Cell Line. International Journal of Molecular Sciences 26, (2025).

56) Seabold, S. & Perktold, J. Statsmodels: Econometric and Statistical Modeling with Python. SciPy 2010 10.25080/Majora-92bf1922-011 (2010) doi:10.25080/Majora-92bf1922-011.

57) Sherry, S. T. et al. dbSNP: the NCBI database of genetic variation. Nucleic Acids Res 29, 308–311 (2001).

58) Silvestris, D. A., Scopa, C., Hanchi, S., Locatelli, F. & Gallo, A. De Novo A-to-I RNA Editing Discovery in lncRNA. Cancers (Basel) 12, 2959 (2020).

59) Smit AFA, Hubley R, Green P. RepeatMasker Open-4.0. 2013-2015. Available from: http://www.repeatmasker.org.

60) Sotgia, F. et al. Mitochondrial metabolism in cancer metastasis: visualizing tumor cell mitochondria and the ‘reverse Warburg effect’ in positive lymph node tissue. Cell Cycle 11, 1445–1454 (2012).

61) Szcześniak, M. W. & Makałowska, I. lncRNA-RNA Interactions across the Human Transcriptome. PLoS One 11, e0150353 (2016).

62) Tan, M. H. et al. Dynamic landscape and regulation of RNA editing in mammals. Nature 550, 249–254 (2017).

63) Tang, S. J. et al. Cis- and trans-regulations of pre-mRNA splicing by RNA editing enzymes influence cancer development. Nat Commun 11, 799 (2020).

64) Virtanen P, Gommers R, Oliphant TE, Haberland M, Reddy T, Cournapeau D, et al. SciPy 1.0: fundamental algorithms for scientific computing in Python. Nat Methods. 2020;17(3):261–272.

65) Vlachogiannis, N. I. et al. Adenosine-to-inosine Alu RNA editing controls the stability of the pro-inflammatory long noncoding RNA NEAT1 in atherosclerotic cardiovascular disease. J Mol Cell Cardiol 160, 111–120 (2021).

66) Walkley, C. R. & Li, J. B. Rewriting the transcriptome: adenosine-to-inosine RNA editing by ADARs. Genome Biol 18, 205 (2017).

67) Wang, I. X. et al. ADAR regulates RNA editing, transcript stability, and gene expression. Cell Rep 5, 849–860 (2013).

68) Zhang, F., Lu, Y., Yan, S., Xing, Q. & Tian, W. SPRINT: an SNP-free toolkit for identifying RNA editing sites. Bioinformatics 33, 3538–3548 (2017).

69) Zhang, Y. et al. Advances in A-to-I RNA editing in cancer. Mol Cancer 23, 280 (2024).

