## Supplementary figures and images for "Antisense lncRNA transcription promotes A-to-I RNA editing via intermolecular dsRNA in breast cancer"

### Supplementary Figure 1

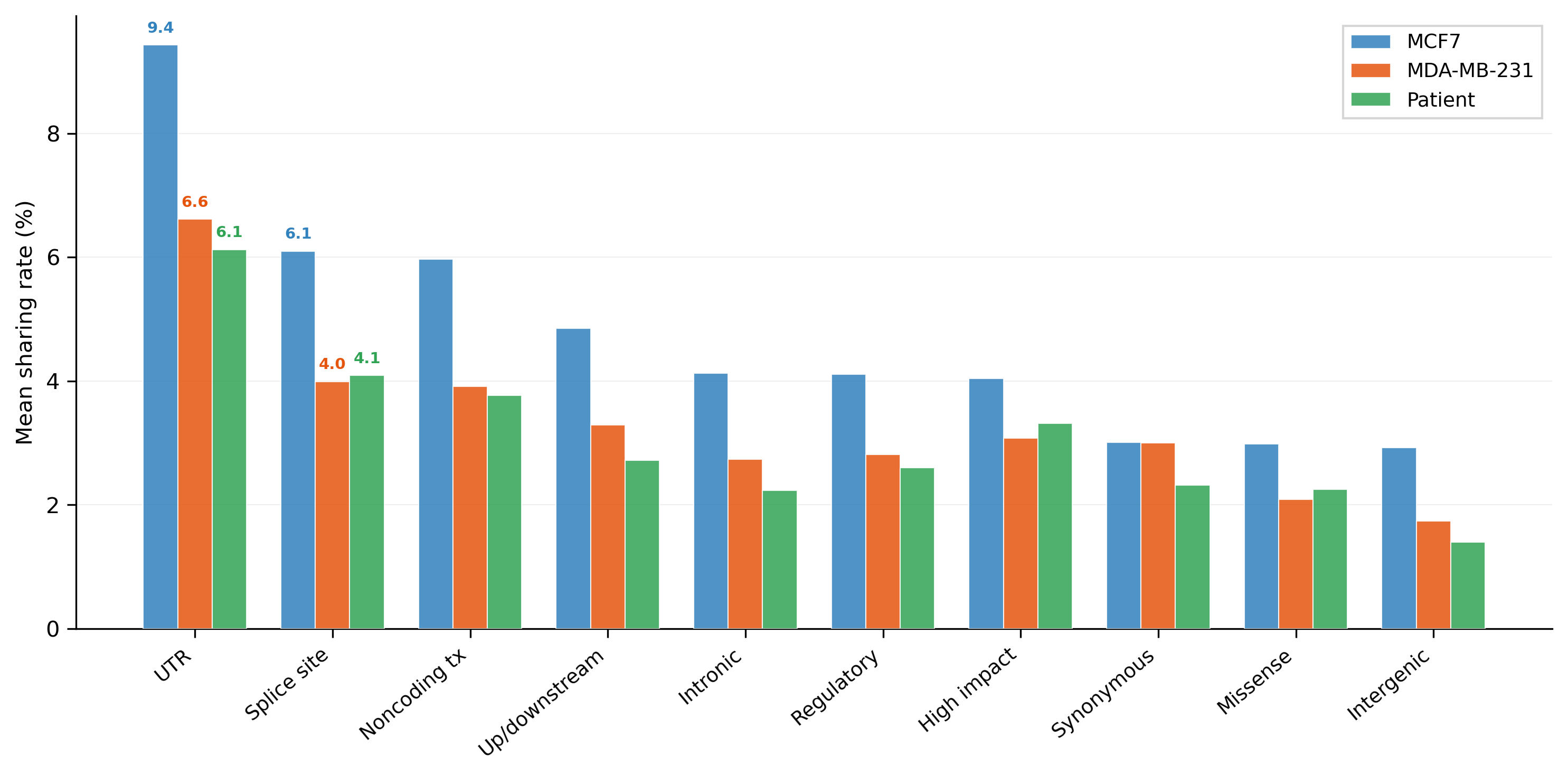

### Supplementary Figure 2

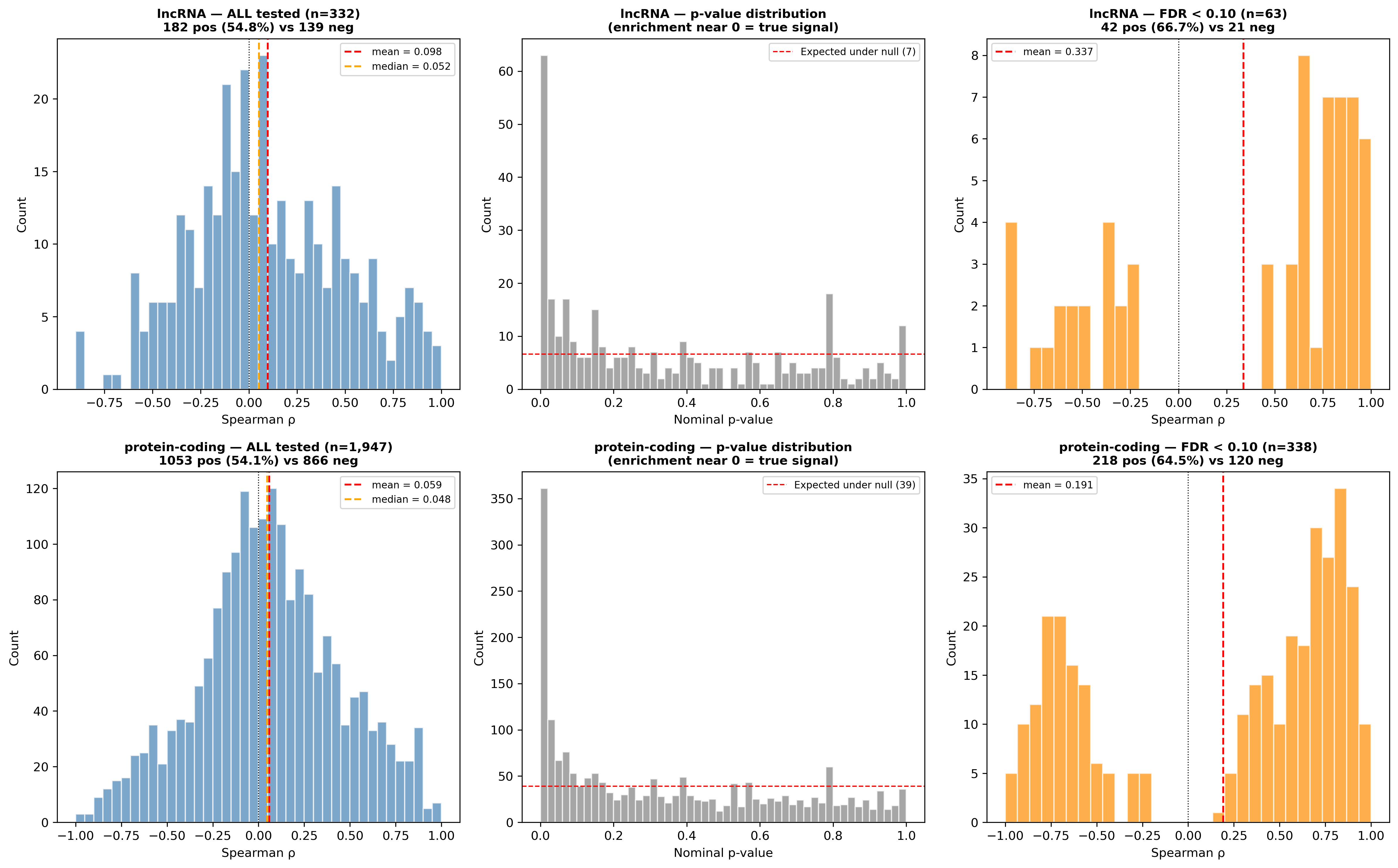

### Supplementary Figure 3

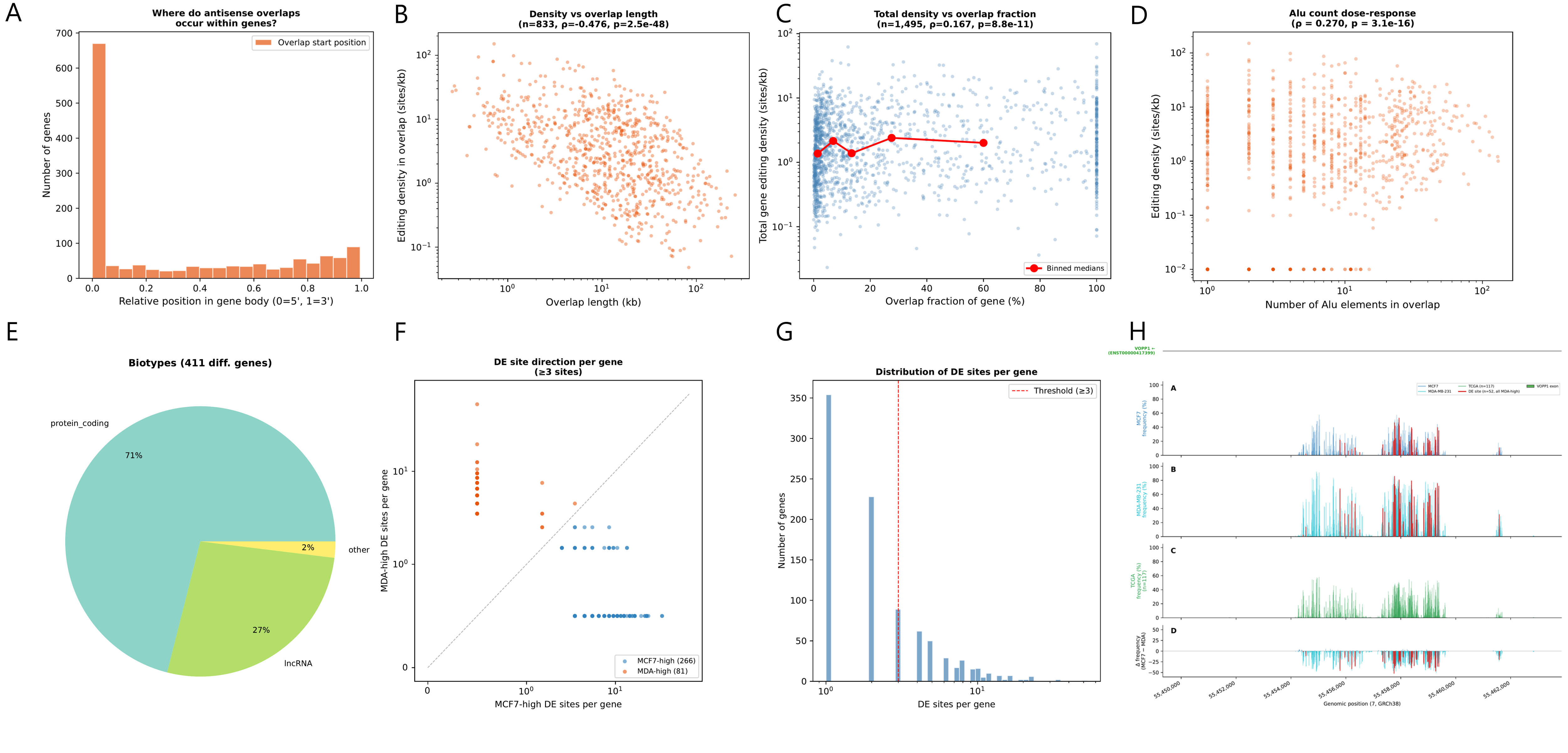

### Supplementary Figure 4

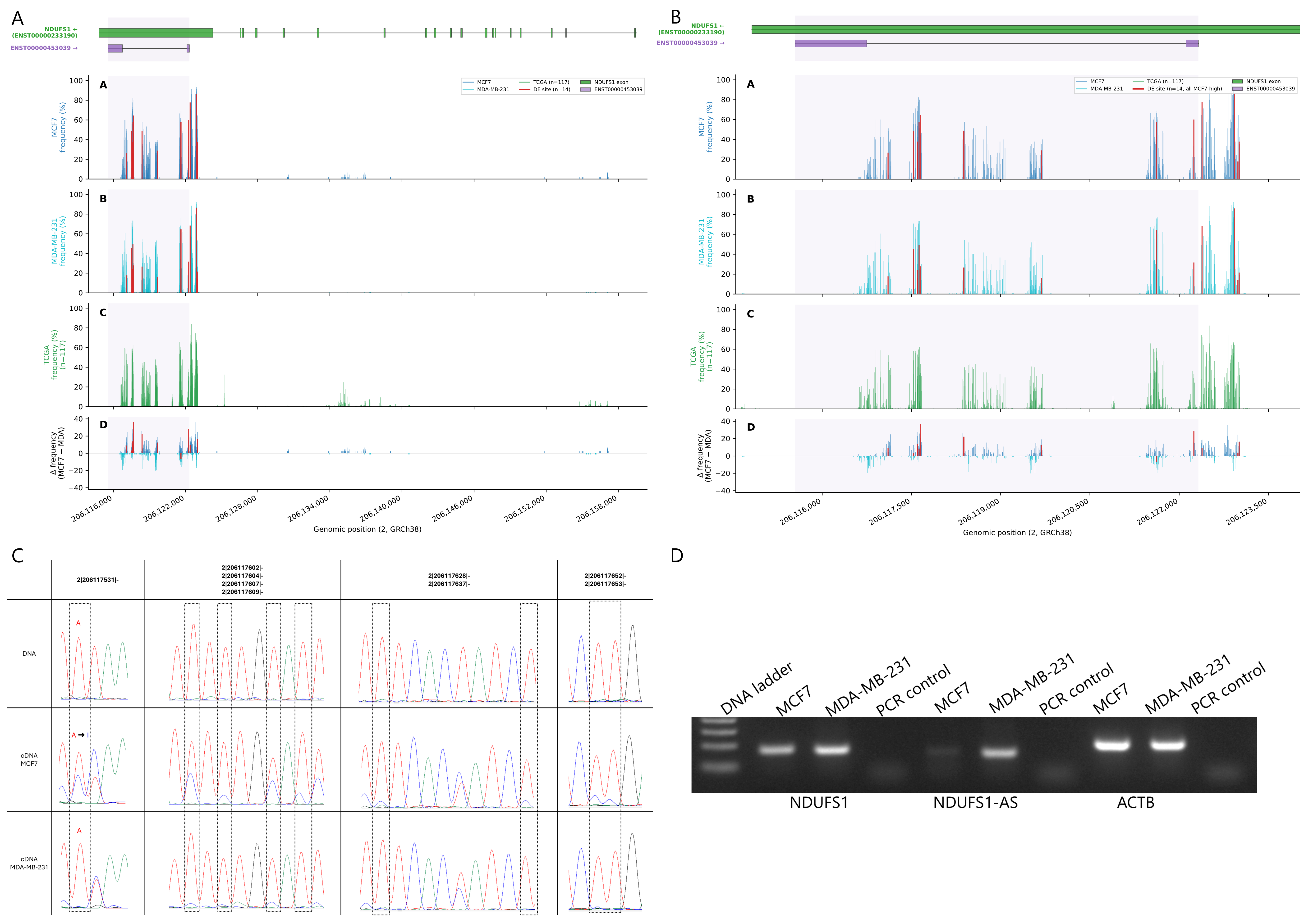
